# Steroid Receptor Coactivator-3 is a Key Modulator of Regulatory T Cell-Mediated Tumor Evasion

**DOI:** 10.1101/2023.03.28.534575

**Authors:** Sang Jun Han, Prashi Jain, Yosef Gilad, Yan Xia, Nuri Sung, Mi Jin Park, Adam M. Dean, Rainer B. Lanz, Jianming Xu, Clifford C. Dacso, David M. Lonard, Bert W. O’Malley

**Affiliations:** Department of Molecular Cellular Biology, Baylor College of Medicine, TX, 77030, USA.; Dan L. Duncan Cancer Center, Baylor College of Medicine, TX, 77030, USA; Department of Medicine, Baylor College of Medicine, TX, 77030, USA

**Keywords:** Adoptive cell transfer, E0771 cell line, Interferon-γ, Regulatory T cells, Steroid Receptor Coactivator 3, Syngeneic murine model of breast cancer

## Abstract

Steroid receptor coactivator 3 (SRC-3) is most strongly expressed in regulatory T cells (Tregs) and B cells, suggesting that it plays an important role in the regulation of Treg function. Using an aggressive E0771 mouse breast cell line syngeneic immune-intact murine model, we observed that breast tumors were ‘permanently eradicated’ in a genetically engineered tamoxifen-inducible Treg-cell specific SRC-3 knockout (KO) female mouse that does not possess a systemic autoimmune pathological phenotype. A similar eradication of tumor was noted in a syngeneic model of prostate cancer. A subsequent injection of additional E0771 cancer cells into these mice showed continued resistance to tumor development without the need for tamoxifen induction to produce additional SRC-3 KO Tregs. SRC-3 KO Tregs were highly proliferative and preferentially infiltrated into breast tumors by activating the Chemokine (C-C motif) ligand (Ccl) 19/Ccl21/ Chemokine (C-C motif) Receptor (Ccr)7 signaling axis, generating antitumor immunity by enhancing the interferon-γ/C-X-C Motif Chemokine Ligand (Cxcl) 9 signaling axis to facilitate the entrance and function of effector T cells and Natural Killer cells. SRC-3 KO Tregs also show a dominant effect by blocking the immune suppressive function of WT Tregs. Importantly, a single adoptive transfer of SRC-3 KO Tregs into wild-type E0771 tumor-bearing mice can completely abolish pre-established breast tumors by generating potent antitumor immunity with a durable effect that prevents tumor reoccurrence. Therefore, treatment with SRC-3 deleted Tregs represents a novel approach to completely block tumor growth and recurrence without the autoimmune side-effects that typically accompany immune checkpoint modulators.

**Significance statement:** Tregs are essential in restraining immune responses for immune homeostasis. SRC-3 is a pleiotropic coactivator, the second-most highly expressed transcriptional coactivator in Tregs, and a suspect in Treg function. The disruption of SRC-3 expression in Tregs leads to a ‘complete lifetime eradication’ of tumors in aggressive syngeneic breast cancer mouse models because deletion of SRC-3 alters the expression of a wide range of key genes involved in efferent and afferent Treg signaling. SRC-3KO Tregs confer this long-lasting protection against cancer recurrence in mice without an apparent systemic autoimmune pathological phenotype. Therefore, treatment with SRC-3 deleted Tregs could represent a novel and efficient future target for eliminating tumor growth and recurrence without the autoimmune side-effects that typically accompany immune checkpoint modulators.

## Introduction

SRC-3 is a pleiotropic coactivator that interacts with a wide range of transcription factors. Cistromic studies of SRC-3 support its pan-genomic transcriptional activities, showing that it is associated with *cis*-regulatory elements across the genome (1). Considering SRC-3’s broad control of gene expression, it is important to consider the mechanistic consequences that are likely to ensue when SRC-3 expression is disrupted. Central to this concept, it is the expectation that perturbation of SRC-3 should result in broad, systemic effects in gene expression and not changes in only a few genes or pathways. Previous studies have shown that whole-body SRC-3 knockout mice show increased B and T cell lymphoproliferation (2). SRC-3 also has a critical role in generating a tumor-promoting immune microenvironment that enhances breast tumor progression in immune-intact mice (3). For example, SRC-3 inhibition with a SRC-3 small molecular inhibitor (SI-2) and SRC-3 knockdown in breast cancer cells generates a tumor-suppressive immune microenvironment in tumors by increasing the numbers of tumor-infiltrating cytotoxic immune cells (such as CD4^+^ and CD8^+^ T cells, NK cells) and Ifng, but reducing CD4^+^Foxp3^+^ regulatory T (Treg) cells compared to controls. Furthermore, SRC-3 is expressed strongly in Tregs (4), suggesting that it may function in Treg-mediated immune suppression. Our observations implicate SRC-3 as a broadly acting and goal oriented pleiotropic regulator of Treg function that can coordinately control the expression of vast numbers of genes in Tregs involved in effecting immune function.

Tregs are essential in restraining immune responses that maintain the immune homeostasis necessary to prevent autoimmune disease. Tregs act by constraining effector immune cell proliferation and function through a number of distinct mechanisms: 1) Tregs secrete cytokines that inactivate effector T cell (CD4+ T and CD8+ T cells) mediated cytotoxicity (5–7). 2) Tregs produce perforin and granzyme B that can induce apoptosis in cytotoxic effector T cells (8). Furthermore, 3) Tregs inhibit NK-cell proliferation and suppress production of interferon-γ (Infg) through their secretion of TGF-β (9). 4) Treg proliferation is strongly induced by exposure to IL-2 secreted by activated T effector cells. These activated and elevated numbers of Tregs produce several anti-inflammatory cytokines (IL-10, IL-35, and TGF-β) that suppress cytotoxic effector immune cells, forming a negative feedback mechanism to limit the immune response (10). Additionally, 5) Tregs interact with monocytes to prevent M1-type macrophage differentiation while enhancing M2-type macrophage differentiation, leading to the production of IL-10 that suppresses effector T cells (11). Notably, the loss of Treg function can result in pathological chronic autoimmune diseases, so a delicate balance must be met to allow the immune system to attack pathogens and cancer cells while also providing an appropriate level of restraint needed to avoid deleterious autoimmune disease (12).

Forkhead box transcription factor 3 (Foxp3) is a principal transcription factor used to define Treg cellular identity (13). While the vast amount of Treg-oriented immuno-therapeutic approaches focus on targeting signaling proteins expressed on the cell membrane, a few studies have sought to understand how transcriptional regulation by Foxp3 modulates Treg-mediated immune homeostasis (14). Focusing on transcriptional regulation in Tregs, we have investigated transcriptional programs within the Treg nucleus, recently identifying SRC-3 as a primary regulator of Treg gene expression (4). SRC-3 has been characterized previously as a prominent oncogene that drives somatic cell cancer progression by activating cell-autonomous growth pathways within cancer cells (15). In addition to cancers, we found previously that SRC-3 has a critical role in immune cell function. For example, SRC-3 is the most highly expressed transcriptional coactivator in Tregs (16). Also, as discussed above, SRC-3 knockout (KO) mice possess an immune phenotype with elevated numbers of B cells and T cells in their lymph nodes, spleen, and bone marrow (2). SRC-3 is highly expressed in both mouse and human Tregs, and inhibition with a SRC-3 small molecular inhibitor (SI-2) inhibits the immune suppressive function of Tregs (4). Therefore, in addition to its function within cancer cells, we suspected that SRC-3 also possesses a critical role within Tregs, and may be responsible for driving immune suppressive functions in Tregs that likely enhance tumor progression.

Herein, we sought to understand the specific role of SRC-3 within the Treg cell compartment, focusing on tumor evasion of the immune system. Using genetically engineered Treg cell-specific SRC-3 KO mice, we show that disruption of Treg SRC-3 expression leads to a ‘complete lifetime eradication’ of tumors in aggressive syngeneic breast and prostate cancer models. Notably, these SRC-3 KO Tregs still support immune checkpoint functions in healthy tissues in the animal, avoiding the severe inflammatory consequences associated with complete loss of Tregs, such as in FoxP3 mutant *scurfy* mice. Notable changes in the expression of Ifng, Il-10, Il-35, and Tgf-β and membrane checkpoint inhibitors were observed in SRC-3 KO Tregs, suggesting that deletion of SRC-3 alters the expression of a range of key genes involved in efferent and afferent Treg signaling, consistent with its role as a known pleiotropic regulator of many mammalian genes. It markedly alters the tumor-immune microenvironment in a way that supports tumor destruction by an animal’s own inherent immune system.

## Results

### Tumors are eradicated in Treg cell-specific SRC-3 knockout mice

To directly determine the role of SRC-3 in Treg cell function, Treg-cell specific SRC-3 KO mice (SRC-^d/d^:Foxp3^Cre/YPF^) were generated by crossing floxed SRC-3 (SRC-3^f/f^, mixed background) mice possessing a flox cassette that brackets known key functional exons 11 and 12 of the *Ncoa3*/*SRC-3* gene mice (17) and Foxp3^Cre/YFP^ mice (B6.129(Cg)-*Foxp3^tm4(YFP/icre)Ayr^*/J. Treg-cell specific SRC-3 KO (SRC-3^d/d^:Foxp3^Cre/YPF^) and WT (SRC-3^f/f^) littermates were generated by breeding SRC-3^d/d^:Foxp3^Cre/YPF^ female mice with SRC-3^f/f^ male mice. The comparative analysis of RNA expression profile in SRC-3 KO Treg versus WT Tregs from spleens of SRC-^d/d^:Foxp3^Cre/YPF^ and SRC-3^f/f^, respectively revealed that SRC-3 and Foxp3 levels were significantly reduced in SRC-3 KO Tregs compared to WT Tregs (Supplement Fig. S1A-C). SRC-3 KO broadly downregulated interleukins, multiple key immune checkpoint genes, chemokine receptors, other key surface proteins, and transcription factors that are required for pro-tumor immune activity in Tregs (Supplement Fig. S1D-F, Supplement Table 1 and 2). Therefore, SRC-3 KO likely would alter the pro-tumor activity of Tregs. To validate our hypothesis, the genetic background of SRC-3^f/f^ mice was moved into the C57BL/6J isogenic line and then crossed with Foxp3-specific tamoxifen-inducible Cre recombinase mice [Foxp3^tm9(EGFP/Cre/ERT2)Ayr^] (Foxp3^Cre-ERT2/+^) to generate bigenic SRC-3^f/f^:Foxp3^Cre-ERT2/+^ mice on a pure C57BL/6J background (Supplement Fig. S2A). SRC-3^f/f^:Foxp3^Cre-ERT2/+^ and SRC-3^f/f^ littermates were generated by breeding SRC-3^f/f^:Foxp3^Cre-ERT2/+^ female mice and SRC-3^f/f^ male mice. After activating the Cre-ERT2 recombinase with tamoxifen in these bigenic SRC-3^f/f^:Foxp3^Cre-ERT2/+^ mice, Treg cell-specific SRC-3 KO (SRC-3^d/d^:Treg mice) were created (Supplement Fig. S2B). Genomic PCR and DNA sequencing of the PCR products validated the deletion of critical functional exons 11 and 12 of the SRC-3 gene in Tregs in spleens of SRC-3^f/f^:Foxp3^Cre-ERT2/+^ female mice (Supplement Fig. S2C and D) (17). Double immunofluorescent staining with antibodies against SRC-3 and Foxp3 in spleens revealed that 65.2% of Foxp3^+^ cells were Foxp3^+^SRC-3^+^ in the spleen of SRC-3^f/f^ female mice, but only 12.1% of Foxp3^+^ cells were Foxp3^+^SRC-3^+^ cells in the spleens of SRC-3^d/d^Treg mice (Supplement Fig. S3A). Therefore, SRC-3 expression was lost in 53.1% of Foxp3^+^ cells in the spleens of SRC-3^d/d^:Treg mice. For further validation of SRC-3 KO in Tregs, Tregs and Tconv isolated from the spleens of SRC-3^f/f^ and SRC-3^d/d^:Treg female mice were analyzed by flow cytometry with an antibody against CD25 and Foxp3 (Supplement Fig. S3B). The percentage of CD25^+^Foxp3^+^ Treg in WT and SRC-3 KO Tregs from the spleens were 80.0 and 92.9 %, respectively. However, the percentage of CD25^+^Foxp3^+^ Tregs counted from total T cells from the spleen of SRC-3^d/d^:Treg was only 18.2%. High purities of isolated WT and SRC-3 KO Treg were achieved. Double immunofluorescent staining revealed that 54.1% of Foxp3^+^ cells were Foxp3^+^SRC-3^+^ in CD4^+^CD25^+^ Tregs from SRC-3^f/f^ mice, but only 13.0% of Foxp3^+^ cells were Foxp3^+^SRC-3^+^ in CD4^+^CD25^+^ Tregs from SRC-3^d/d^:Treg mice (Supplement Fig. S3C). Also, comparison of SRC-3 mRNA levels in Tregs from the spleens of SRC-3^d/d^:Treg female mice with their SRC-3^f/f^ controls revealed a significant reduction in expression (Supplement Fig. S3D). Collectively, these data show that SRC-3 expression is strongly abrogated in Tregs of SRC-3^d/d^:Treg female mice.

Body weights of SRC-3^d/d^:Treg female mice over their life-time was not statistically different from that of SRC-3^f/f^ female mice (Supplement Fig. S4A). Reproductive capacity is a hallmark of mouse health. Notably, mouse fertility analysis revealed that the number of pups produced by SRC-3^d/d^:Treg female mice mated with wild-type male mice was not statistically different from that produced by SRC-3^f/f^ female mice mated with wild-type male mice (Supplement Fig. S4B). *Foxp3*^−/-^ *scurfy* mice possess a severe, systemic autoimmune phenotype and die at an early age (18, 19). Unlike *scurfy* mice, however, SRC-3^d/d^:Treg female mice have similar levels of blood cytokine/chemokines compared to SRC-3^f/f^ female mice, with only three observable cytokines (Timp-1, Ccl2, and Il-1ra) elevated in the SRC-3^d/d^:Treg mice (Supplement Fig. S4C and D). In addition to their cytokine profiles, immuno-phenotyping analysis of the spleens of SRC-3^d/d^:Treg mice did not show an altered repertoire of lymphocytes and myeloid cells compared to that of SRC-3^f/f^ female mice (Supplement Fig. S4E and F). These data indicate that SRC-3^d/d^:Treg mice do not possess the pathological characteristics observed in mice with entirely abrogated Treg function or with checkpoint inhibitor agents.

E0771 cells, an aggressive breast cancer cell line established from a spontaneous mammary gland carcinoma in the C57BL/6 mouse strain, were used to characterize how SRC-3 KO Tregs impact breast cancer progression in immune-intact mouse (20). E0771 cells were orthotopically injected into the mammary fat pads of SRC-3^d/d^:Treg and SRC-3^f/f^ female mice (Supplement Fig. S5A). E0771 breast tumors grew aggressively in SRC-3^f/f^ female mice (Fig. 1A and B). Early E0771 breast tumors were detected in SRC-3^d/d^:Treg female mice five days after E0771 cell injection, but breast tumors became undetectable after 20 days (Fig. 1A and B). Hematoxylin and eosin (H&E) analyses validated eradication of the primary breast tumors in SRC-3^d/d^:Treg female mice and no tumor masses were detected in mammary fat pads of these animals (Fig. 1C). Also, SRC-3^d/d^:Treg female mice did not show splenomegaly, with spleens appearing similar to SRC-3^f/f^ female control mice (Fig. 1D). In Foxp3^Cre-ERT2/+^ monogenic mouse lines, E0771 breast tumors also grew aggressively (Supplement Fig. S6A-D). These observations confirm that the lack of tumor growth in the SRC-3^f/f^:Foxp3^Cre-ERT2/+^ bigenic mice was not due to mouse background genetic issues. To rule out the potential direct effects of tamoxifen on breast tumors, tumor-bearing SRC-3^f/f^ female mice were treated with tamoxifen versus vehicle (Supplement Fig. S7A). Tumor luciferase activity analysis revealed that tamoxifen treatment did not impact the growth of E0771 breast tumors in SRC-3^f/f^ female control mice (Supplement Figs. S7B and C).

**Fig. 1.**
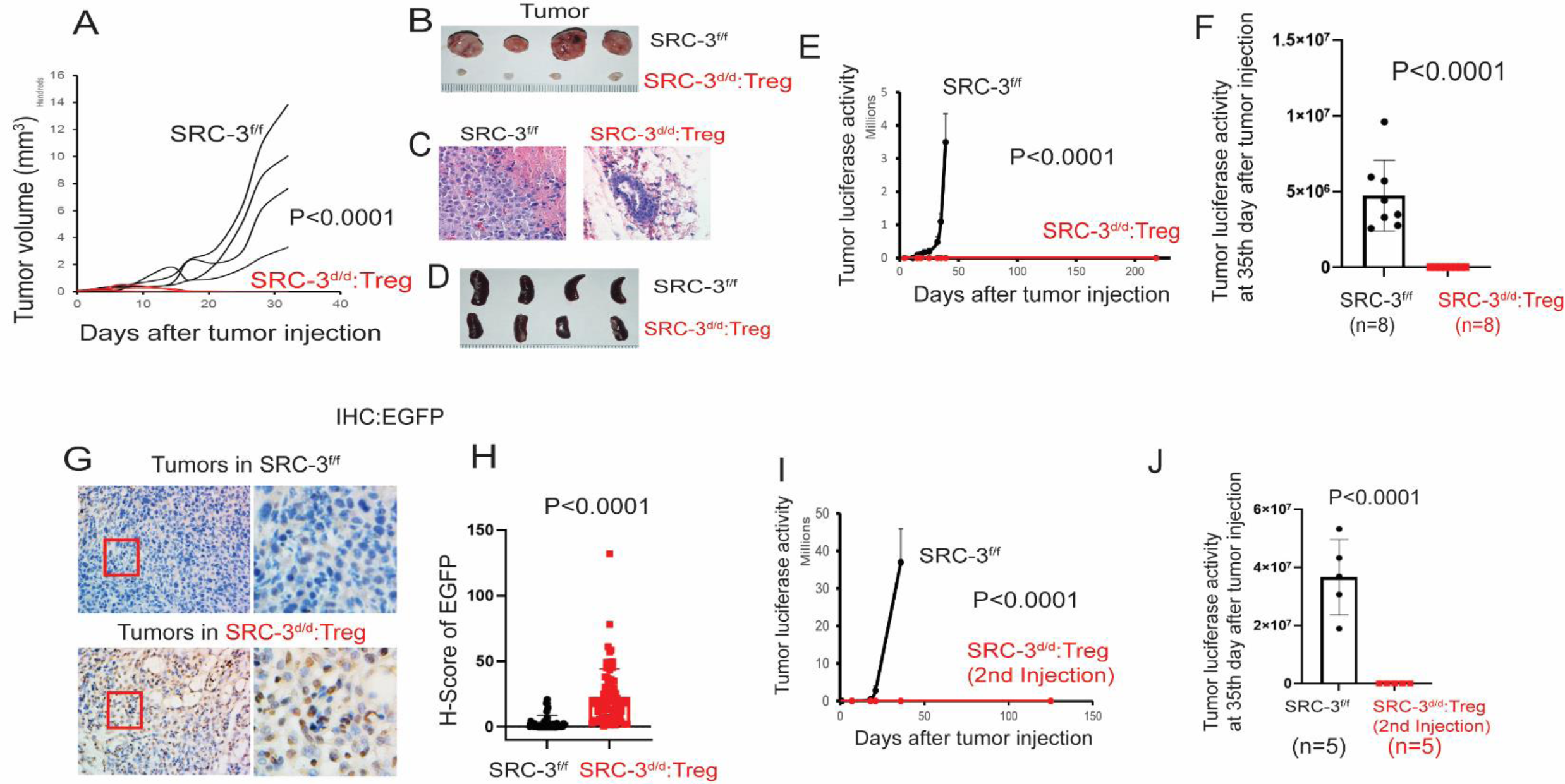
**E0771 breast tumor eradication in SRC-3^d/d^:Treg mice.** (A) E0771 breast tumor growth in SRC-3^f/f^ versus SRC-3^d/d^:Teg female mice. (B) Breast tumors isolated from SRC-3^f/f^ and SRC-3^d/d^:Teg female mice on the 33^rd^ day after injection with E0771 cells. (C) H&E staining of tumors is presented in panel (B). (D) Spleens isolated from SRC-3^f/f^ and SRC-3^d/d^:Teg female mice on the 40^th^ day after tamoxifen treatment. (E) Breast tumor luciferase activity in SRC-3^f/f^ and SRC-3^d/d^:Treg female mice. (F) The repeated tumor luciferase activity experiment with tamoxifen-treated SRC-3^f/f^ (n=8) and SRC-3^d/d^:Treg mice (n=8) was determined on the 35^th^ day after E0771:LUC cell injection. (G) IHC analysis of EGFP+ cells in tumors of tamoxifen-treated SRC-3^f/f^ and SRC-3^d/d^:Treg mice on the 14^th^ day after E0771 cell injection. (H) H-score for EGFP+ cells in tumors presented in panel (G). (I) Tumor luciferase activity in SRC-3^f/f^ and breast cancer-eradicated SRC-3^d/d^:Treg mice after a 2^nd^ injection of E0771:LUC cells. (J) The repeated experiment for tumor luciferase activities in tamoxifen-treated SRC-3^f/f^ (n=5) and breast tumor-eradicated SRC-3^d/d^:Treg mice (n=5) on the 35^th^ day after E0771:LUC cell reinjection.

To noninvasively analyze tumor progression, stable luciferase-expressing E0771 (E0771:LUC) cells were developed by infection with a lentiviral luciferase expression vector. To determine whether one injection of SRC-3 KO Tregs can prevent tumor initiation, E0771:LUC cells were injected into SRC-3^f/f^:Foxp3^Cre-ERT2/+^ female mice after first treating them with tamoxifen or vehicle control (Supplement Fig. S8A). E0771 breast tumors developed aggressively in SRC-3^f/f^:Foxp3^Cre-ERT2/+^ female mice treated with vehicle, but tumors failed to progress beyond a short initial period of growth in SRC-3^f/f^:Foxp3^Cre-ERT2/+^ female mice treated with tamoxifen (Supplement Fig. S8B and C). Next, we sought to examine further whether SRC-3 KO Tregs can drive the regression of ‘pre-existing’ E0771 tumors in mice. SRC-3^f/f^:Foxp3^Cre-ERT2/+^ and SRC-3^f/f^ female mice were first injected with E0771:LUC breast cancer cells. After tumors were allowed to grow for seven days, mice were treated with tamoxifen (Supplement Fig. S8D). Tumors again were eradicated in SRC-3^f/f^:Foxp3^Cre-ERT2/+^ mice that were treated with tamoxifen. In contrast, tumors continued to progress in tamoxifen-treated SRC-3^f/f^ female mice (Supplement Fig. S8E and F). Compared to tamoxifen treatment, breast tumors grew aggressively in SRC-3^f/f^:Foxp3^Cre-ERT2/+^ female mice treated with vehicle (Supplement Fig. S8G, H, and I). Therefore, SRC-3 KO Tregs induced with tamoxifen treatment eradicate breast tumors in SRC-3^d/d^:Treg female mice.

The biggest problem in current cancer therapies is tumor recurrence after cessation of treatment. Thus, breast tumor recurrence in SRC-3^d/d^:Treg female mice was examined. E0771:LUC cells were orthotopically injected into SRC-3^f/f^ and SRC-3^d/d^:Treg female mice. E0771 breast tumors became undetectable in SRC-3 ^d/d^:Treg female mice 20 days after E0771:Luc cell injection, and tumor recurrence was not detected in SRC-3^d/d^:Treg female mice for >218 days after E0771:LUC cell injection (Fig. 1E and Supplement Figs. S9A and B); we consider these tumors to be permanently eliminated from the SRC-3^d/d^:Treg female mice. We repeated this experiment with increased animal numbers, confirming the complete eradication of tumors in SRC-3 ^d/d^:Treg female mice (n=8), but not in SRC-3^f/f^ female mice (n=8) (Fig. 1F).

The SRC-3 KO Tregs express enhanced-EGFP as a result of the bigenic animals possessing the Foxp3^tm9(EGFP/Cre/ERT2)Ayr^ cassette (Supplement Fig. S2B). Tumors were isolated from SRC-3^d/d^:Treg and SRC-3^f/f^ female mice 14 days after E0771 cell injection (at an early time point when tumors were not yet entirely eradicated in SRC-3^d/d^:Treg female mice). IHC analysis with an EGFP antibody revealed the presence of large numbers of EGFP+ Tregs within the tumors of SRC-3^d/d^:Treg female mice but not in tumors of SRC-3^f/f^ control mice (Fig. 1G and H). These results indicate that SRC-3 KO Tregs infiltrate into E0771 tumors before tumor eradication in the SRC-3^d/d^:Treg mice.

The above observation raised the question of whether another type of cancer can be eradicated in the SRC-3^d/d^:Treg mouse model. To address this question, the RM-1 syngeneic immune-intact mouse model of prostate cancer was employed (21). Orthotopically injected luciferase-labeled RM-1 prostate cancer cells developed into prostate cancers in SRC-3^f/f^ male mice but not in SRC-3^d/d^:Treg male mice (Supplement Fig. S10A). Furthermore, after prostate tumors disappeared, prostate cancer recurrence was not detected in the SRC-3^d/d^:Treg males extending to at least >150 days (Supplement Fig. S10B). Thus in a similar fashion to that observed in the E0771 breast cancer model, we found that SRC-3 KO Tregs also can eliminate prostate tumors in an aggressive male prostate cancer mouse model.

The lack of E0771 tumor recurrence in SRC-3^d/d^:Treg mice raised the question of whether long-term resistance to subsequent new tumor formation would occur. To address this question, E0771:LUC cells were reinjected into tumor-eradicated SRC-3^d/d^:Treg mice (Supplement Fig. S5B) and SRC-3^f/f^ female mice as controls. Tumors grew rapidly in SRC-3^f/f^ control female mice but ‘reinjected’ E0771 cells could not establish breast tumors in tumor-eradicated SRC-3^d/d^:Treg female mice (at least >125 days) (Fig. 1I and Supplement Fig. S9C). A repeated experiment with an increased number of animals confirmed the tumor resistance phenotype in all SRC-3^d/d^:Treg female mice (n=5) (Fig. 1J).

Next, we wished to determine how SRC-3 KO Tregs support tumor eradication at a cell mechanistic level with a focus on how these cells modulate other immune cells in the tumor microenvironment. Since E0771 breast tumors were still present and available for analysis in SRC-3^d/d^:Treg female mice until 14 days after E0771 cell injection (Fig. 2A), tumors were isolated from these mice and SRC-3^f/f^ control mice at this time. H&E staining revealed extensive hypocellularity in tumors from the SRC-3^d/d^:Treg mice but not in SRC-3^f/f^ mice (Fig. 2B). Tregs usually suppress and inactivate effector immune cells including CD4^+^ T, CD8^+^ T, and natural killer (NK) cells in the tumor immune microenvironment, facilitating tumor progression. The numbers of infiltrating CD4^+^ T cells in tumors from SRC-3^d/d^:Treg mice when compared to that from SRC-3^f/f^ mice (Fig. 2C), CD8^+^ T cells and CD49b^+^ NK cells were significantly elevated in tumors from SRC-3^d/d^:Treg female mice when compared to SRC-3^f/f^ mice (Figs. 2D and E). Foxp3 immunostaining revealed fewer Foxp3+ Tregs in tumors from SRC-3^d/d^:Treg mice compared to SRC-3^f/f^ control mice (Fig, 2F). Therefore, the markedly increased number of infiltrating CD8^+^ and NK effector immune cells observed in tumors from the SRC-3^d/d^:Treg mice was consistent with establishment of an antitumor immune microenvironment by the SRC-3 KO Tregs. To determine the Treg population in tumors, Treg cells were isolated from tumors of tumor-bearing SRC-3^f/f^ and SRC-3^d/d^:Treg female mice, as described above. Flow cytometry analysis of antibodies against Foxp3 and SRC-3 showed that the percentage of Foxp3^+^SRC-3^−^ population in total tumor Foxp3^+^ Tregs in SRC-3^f/f^ and SRC-3^d/d^:Treg female mice was 1.6% and 17.4%, respectively (Supplement Fig. S12A and B). Interestingly, Foxp3^−^SRC-3^−^ Tregs were significantly increased in tumors in SRC-3^d/d^:Treg female mice (25.6%) compared to tumors in SRC-3 ^f/f^ female mice (6.3%). In the spleen, the percentage of Foxp3^+^SRC-3^−^ Tregs out of total Foxp3^+^ Tregs in SRC-3^f/f^ and SRC-3^d/d^:Treg female mice was 3.9% and 41.1%, respectively (Supplement Fig. S12C and D). However, Foxp3^−^SRC-3^−^ Tregs were not elevated in the spleens of SRC-3^d/d^:Treg female mice (3.85%) compared to SRC-3 ^f/f^ female mice (2.37%). Therefore, the elevation of both Foxp3^+^SRC-3^−^ and Foxp3^−^SRC-3^−^ Tregs can generate a tumor-suppressive immune microenvironment in SRC-3^d/d^:Treg female mice sufficient for breast tumor eradication.

**Fig. 2.**
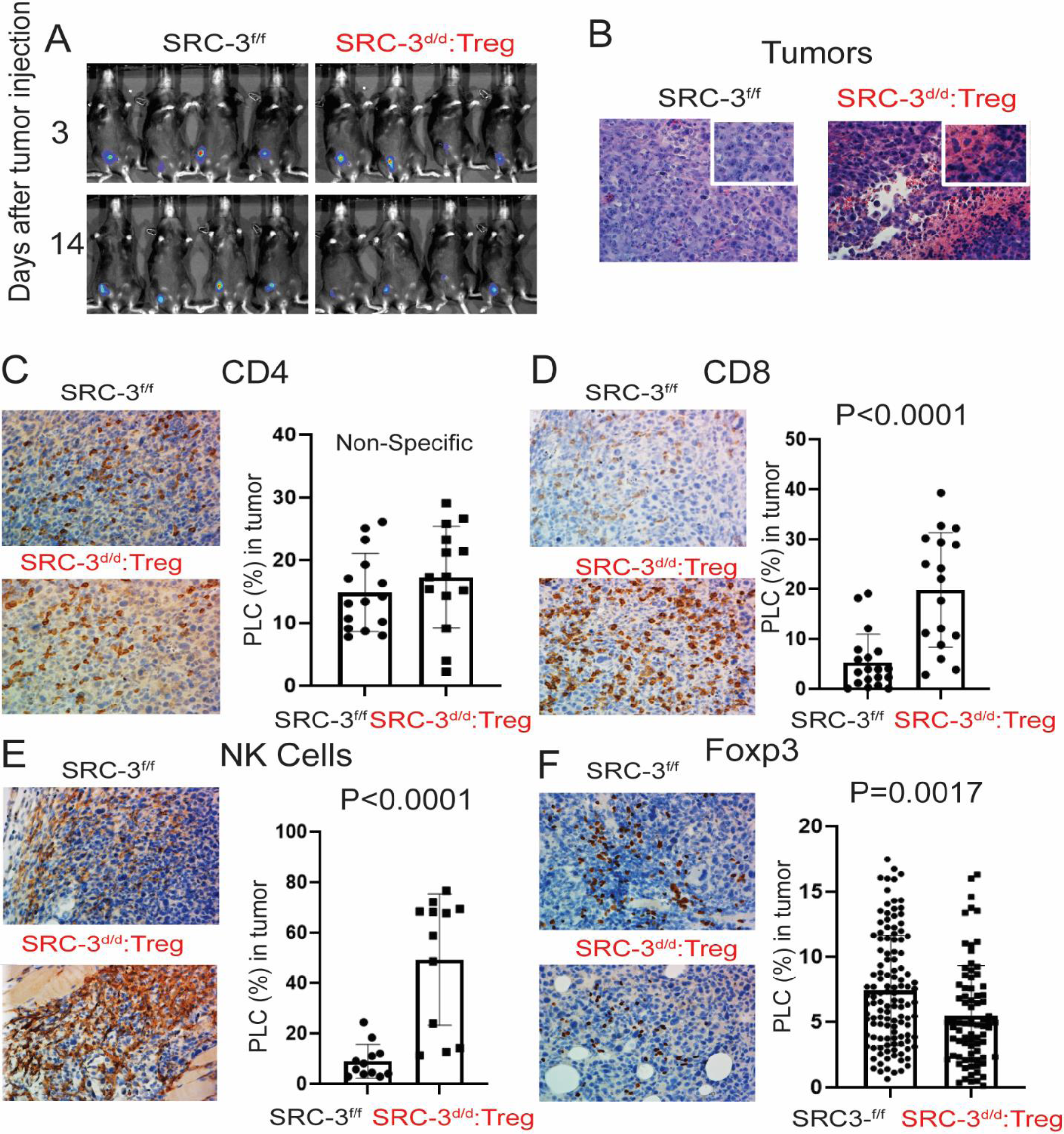
SRC-3^d/d^:Treg mice generate a tumor suppressive immune microenvironment in breast tumors. (A) Images of tumor luciferase activity in SRC-3^f/f^ and SRC-3^d/d^:Treg mice on the 3^rd^ and 14^th^ days after E0771:LUC cell injection. (B) H&E staining of tumors in SRC-3^f/f^ and SRC-3^d/d^:Treg mice on the 14^th^ day after E0771:LUC cell injection. (C-F) IHC analysis of CD4^+^ T cells (C), CD8^+^ T cells (D), CD49b^+^ NK cells (E), and Foxp3+ Tregs (F) in tumors from SRC-3^f/f^ and SRC-3^d/d^:Treg mice on the 14^th^ day after E0771:LUC cell injection. The QuPath program was used to quantify IHC staining results. PLC, Percentage of Labelled Cells.

The cytokine/chemokine milieu within tumors alters the tumor immune microenvironment and strongly impacts cancer progression (22). To investigate the consequences of SRC-3 KO Treg infiltration into E0771 tumors, we performed cytokine/chemokine profiling from the tumors of SRC-3^d/d^:Treg and SRC-3^f/f^ mice 14 days after E0771:LUC cell injection; we observed that levels of Ifng, Cxcl9 and Mip-1α are significantly elevated in tumors of SRC-3^d/d^:Treg female mice compared to controls (Fig. 3A and B). In addition, immunohistochemical (IHC) analysis validated marked increases in Ifng and Cxcl9 levels in tumors from SRC-3^d/d^:Treg compared to SRC-3^f/f^ control mice (Fig. 3C and D). To determine whether Ifng is a critical factor in breast cancer eradication by SRC-3 KO Tregs, breast tumor-bearing SRC-3^f/f^ and SRC-3^d/d^:Treg female mice were intraperitoneally injected with anti-Ifng antibody and rat IgG as the control IgG (Supplement Fig. S12A). The Ifng depletion by anti-Ifng antibody reactivated E0771 breast tumor progression in SRC-3^d/d^:Treg female mice compared to control IgG (Fig. 3E, Supplement Fig. S12B). However, anti-Ifng antibody treatment did not impact the breast tumor growth in SRC-3^f/f^ female mice (Fig. 3F, Supplement Fig. S12B). Therefore, Ifng appears to be essential for breast tumor eradication in SRC-3^d/d^:Treg female mice. Ifng is a central proinflammatory cytokine that can suppress tumor progression and activate CD8^+^ T cells and NK cells (23, 24). Cxcl9 is a chemokine that recruits and activates Cxcr3^+^ immune cells, including CD8^+^ and NK cells (25). We conclude that SRC-3 KO Tregs strongly enhance the Ifng/Cxcl9 axis in E0771 tumors to support the establishment of an antitumor immune microenvironment by recruiting and activating CD8^+^ and NK effector immune cells. In contrast to its large increase in the tumor microenvironment, Ifng was <u>not</u> systemically elevated in serum from these animals (Supplement Fig. S4C).

**Fig. 3.**
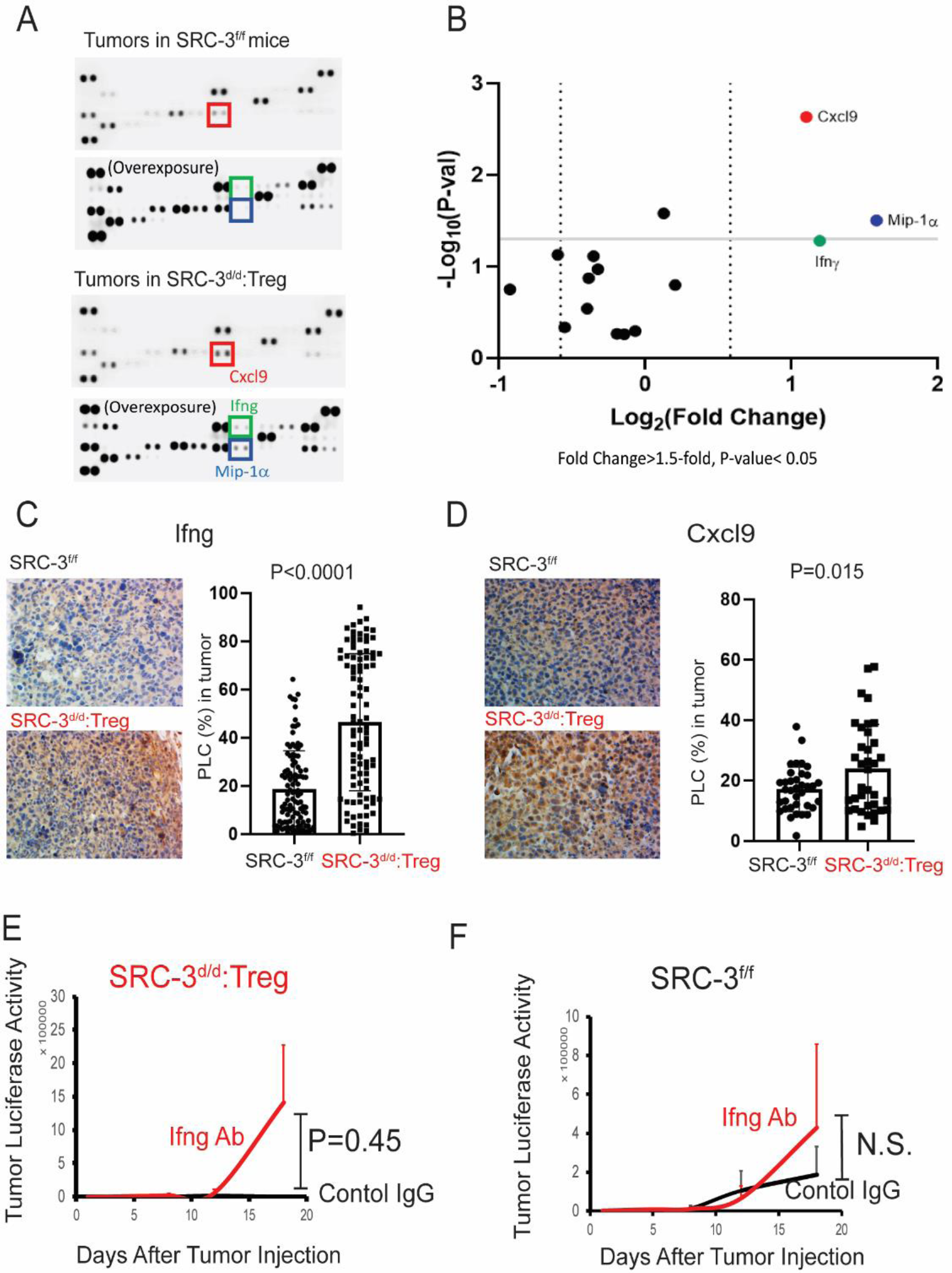
SRC-3^d/d^:**Treg mice possess an elevatd Ifng/Cxcl9 signaling axis in breast tumors.** (A) Cytokine/chemokine profiling in breast tumors obtained from SRC-3^f/f^ and SRC-3^d/d^:Treg female mice on the 14^th^ day after E0771:LUC cell injection. (B) Quantification of cytokines and chemokines shown in panel (A). (C and D) IHC analysis of Ifng (C), and Cxcl9 (D) in tumors from SRC-3^f/f^ and SRC-3^d/d^:Treg mice on the 14^th^ day after E0771:LUC breast cancer cell injection using the QuPath program. (E-F) Tumor luciferase activity of tumor-bearing SRC-3^d/d^:Treg (E) and SRC-3^f/f^ (F) female mice treated with anti-Ifng antibody (Ifng Ab) and control Rat IgG (Control IgG). PLC, Percentage of Labelled Cells.

### SRC3 KO Tregs possess a ‘functional’ dominant role in tumor immunity

How doSRC-3 KO Tregs eradicate breast tumors even though WT Tregs exist in the animals and in breast tumors? Treg RNA analysis revealed the downregulation of Tigit, Klrb1c, and Klrk1 in SRC-3 KO Tregs compared to WT Tregs (Fig. 4A-C and Supplement Fig. S1F). Since Tigit, Klrb1c, and Klrk1 are co-inhibitory molecules in Tregs that inhibit pro-inflammatory T cell responses selectively (26), the downregulation of these co-inhibitory molecules after SRC-3 KO would be expected to disrupt immune suppressive functions of Tregs. Tregs also were isolated from tumors and spleens of tumor-bearing SRC-3^d/d^:Treg and SRC-3^f/f^ mice 14 days after E0771 breast cancer cell injection. RNA analysis by quantitative PCR (qPCR) with Treg cell marker genes revealed that SRC-3 KO Tregs in spleens of tumor-bearing SRC-3^d/d^:Treg female mice had elevated expression of anti-inflammatory cytokines such as Tgf-β, Il-10, and Il-35 compared to Tregs from the spleens of tumor-bearing SRC-3^f/f^ female mice (Fig. 4D). In contrast, SRC-3 KO Tregs in tumors from tumor-bearing SRC-3^d/d^:Treg female mice had significantly elevated expression of Ifng (∼5-fold) compared with Tregs from the tumors in tumor-bearing SRC-3^f/f^ female mice (Fig. 4E). These results reveal that the molecular phenotypes of SRC-3 KO Tregs are very different in spleens versus tumors, and SRC-3 KO Tregs in breast tumors promotes the antitumor environment by inducing expression of Ifng and other cytokines. Thus, T Cell Receptor (TCR) activation appears likely to be a critical event for the induction of Ifng in SRC-3 KO Tregs in breast tumors.

**Fig. 4.**
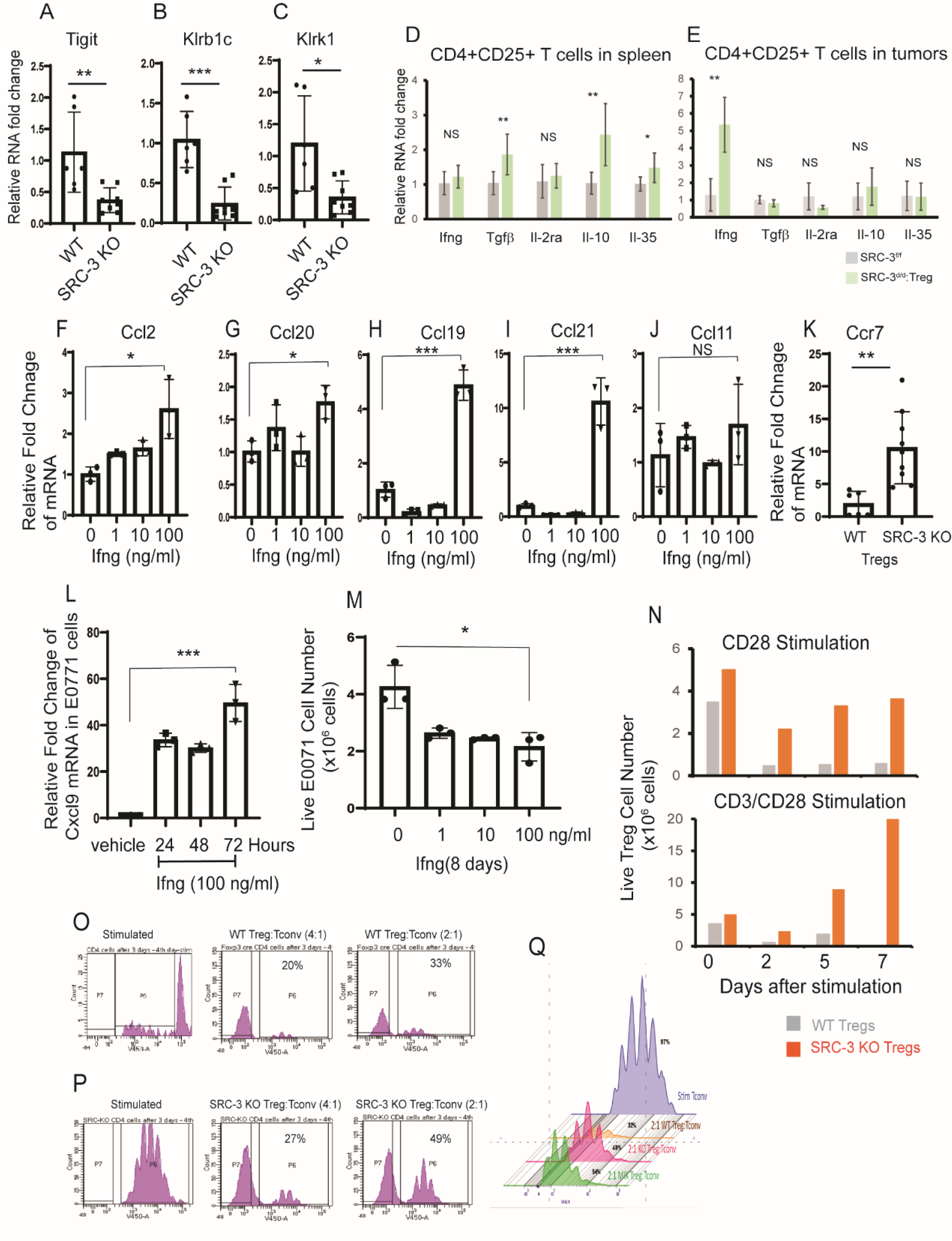
Molecular properties of SRC-3 KO Treg cells. (A-C) Relative RNA levels of Tigit (A), Klrb1c (B), and Klrk1 (C) in WT versus SRC-3 KO Treg cells from spleens. (D and E) Relative mRNA levels for Ifng, Tgf-β, Il-2ra, Il-10, and Il-35 in CD4^+^CD25^+^ Treg cells isolated from spleens (D) and breast tumors (E) in breast tumor-bearing SRC-3^f/f^ and SRC-3^d/d^:Treg mice on the 14^th^ day after E0771:LUC cell injection. (F-J) Relative RNA levels of Ccl2 (F), Ccl20 (G), Ccl19(H), Ccl21(I), and Ccl11(J) in E0771 breast cancer cells treated with 0, 1, 10, and 100 ng/ml of Ifng. (K) RNA levels of Ccr7 in WT versus and SRC-3 KO Tregs. (L) RNA analysis of Cxcl9 in E0771 breast cancer cells upon Ifng (100 ng/ml) treatment for 24, 48, and 72 hours. (M) The viability of E7001 breast cell numbers upon 1, 10, and 100 ng/ml of Ifng for 8 days. (N) The viability of WT Tregs and SRC-3 KO Tregs at 0, 2, 5, and 7^th^-day after CD28 and CD3/CD28 stimulation. (O) Flow cytometry analysis of T cells showing immune suppressive activity of WT Tregs. (P) Flow cytometry analysis of T cells showing the reduced immune suppressive activity of SRC-3 KO Tregs. (Q) The reduced suppressive activity of the mixture of SRC-3 KO and WT Tregs (1:1) to Tconv cell proliferation when compared to WT Tregs. *, p<0.05; **, P<0.01; ***, p<0.001; NS, nonspecific

What is the role of Infg in breast tumor eradication by SRC-3 KO Tregs? Ifng treatment significantly increased Ccl19/Ccl21 (Treg chemotactic ligands) in E0771 breast cancer cells in contrast to other Treg chemotactic ligands such as Ccl2, Ccl20 and Ccl 11 (Fig. 4 F-J). Our RNA-seq analysis revealed that the level of Ccr7 (Ccl19/Ccl21 receptor) was significantly elevated in SRC-3 KO Tregs compared to WT Tregs (Fig. 4K). Therefore, Ifng activates the Ccl19/Ccl21/Ccr7 axis between breast tumors and SRC-3 KO Tregs, to signal the recruitment of SRC-3 KO Tregs into breast tumors. Ifng treatment also significantly increased the level of Cxcl9 mRNA expression in E0771 cells (Fig. 4L) and suppressed the viability of E0771 cells compared to vehicle (Fig. 4M). Thus, Ifng from SRC-3 KO Tregs elevates Cxcl9 expression in E0771 breast tumors in SRC-3^d/d^:Treg female mice and suppresses the growth of breast tumors. Additional *in vitro* cell survival analysis with Tregs shows that the viability of WT Treg cells from spleens of SRC-3^f/f^ female mice declined over a seven day period, but the viability of SRC-3 KO Tregs was not reduced over the same time period (Fig, 4N). Also, in the presence of CD3/CD28 co-stimulation, the viability and proliferation of SRC-3 KO Tregs was significantly elevated during *in vivo* culture compared to WT Tregs (Fig. 4N). This data is consistent with our findings that SRC-3 KO Tregs are hearty cells that ‘functionally predominate’ over WT Tregs in the tumor microenvironment.

Next, we sought to determine whether SRC-3 KO impacts the immune suppressive function of Tregs using co-culture assays consisting of CD4^+^ CD25^+^ (Tregs) mixed with conventional CD4^+^ CD25^−^ (Tconv) cells from the spleens of SRC-3^d/d^:Treg and SRC-3^f/f^ female mice. Wild-type Tregs suppressed stimulated Tconv cell proliferation (Fig. 4O), while SRC-3 KO Tregs suppressed Tconv cell proliferation to a substantially smaller extent (Fig. 4P). Moreover, a 2:1 mixture of wild-type and SRC-3 KO Tregs led to the same attenuated suppression of Tconv cell proliferation (Fig. 4Q). Therefore, SRC-3 KO Tregs effectively inhibited the Tconv suppressive activity of WT Tregs. Collectively, the above observations suggest that SRC-3 KO Tregs have a ‘functionally dominant’ role in eradicating breast cancer, irrespective of the suppressive effects of neighboring wild-type Tregs.

### Adoptive cell transfer (ACT) with SRC-3 KO Tregs eliminates E0771 tumors

ACT with Tregs has been promoted as an immunotherapy to treat various chronic autoimmune diseases, but not for treating solid cancers (27). Since SRC-3 KO Tregs have a tumor-eradicating activity and a suppressive activity against WT Tregs, we wished to determine whether a ‘single injection’ of ACT with purified SRC-3 KO Tregs would eradicate pre-existing E0771 tumors growing in wild-type mice by suppressing the inherent function of WT Tregs. SRC-3 KO and control wild-type Tregs were isolated from the spleens of SRC-3^d/d^:Treg and SRC-3^f/f^ female mice, respectively, and then injected Tregs (800K cells) into tumor-bearing SRC-3^f/f^ female littermates (Supplement Fig. S13A and B). E0771:LUC cells were developed into breast tumors in SRC-3^f/f^ female mice before conducting ACT (Supplement Fig. S14A). Notably, ACT with SRC-3 KO Tregs; completely eradicated’ established tumors in SRC-3^f/f^ littermates, but ACT with purified wild-type Tregs or no ACT did not suppress breast tumor growth in SRC-3^f/f^ littermates (Fig. 5A and Supplement Fig. S14B). Furthermore, after the initial regression of tumors to undetectable levels through ACT with SRC-3 KO Tregs, tumor recurrence was not observed in these animals for >215 days after ACT (Fig. 5A).

**Fig 5.**
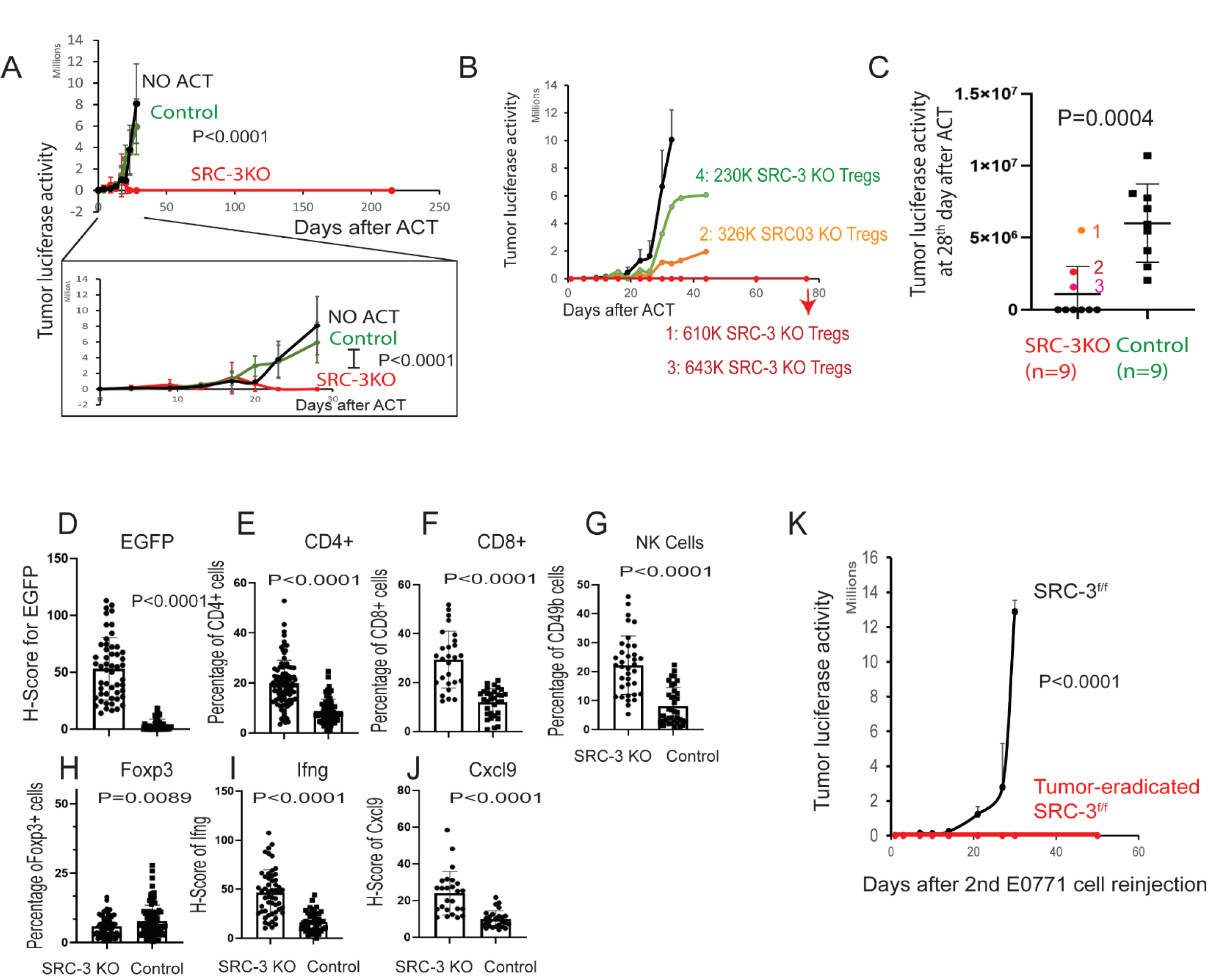
Tumor eradication through adoptive SRC-3 KO Treg transfer. (A) Tumor luciferase image analysis in tumor-bearing SRC-3^f/f^ female mice after ACT with SRC-3 KO Tregs (800K cells) and wild-type Tregs (800K cells). Control tumor-bearing SRC-3^f/f^ mice that did not undergo ACT (NO ACT). Tumor luciferase image analysis up to the 30^th^ day, which is in the box, and 215^th^ days after ACT. (B) Tumor luciferase activity in SRC-3^f/f^ female mice after ACT with different doses of SRC-3 KO Tregs. Tumor-bearing SRC-3^f/f^ female mice were adoptively transferred with wild-type (800K cells) or SRC-3 KO Treg cells (1: 610K cells, 2: 326K cells, 3: 643K cells, 4: 230K cells). Afterward, the luciferase activity emanating from tumors was determined. (C) Repeated ACT experiment with increase animal numbers. Tumor luciferase activity in SRC-3^f/f^ female mice adoptively transferred with SRC-3 KO Tregs (1: 230K cells, 2: 314K cells, 3: 358K cells, others: 800K cells, n=9) and wild-type Tregs (800K cells, n=9) was determined on the 28^th^ day after E0771:LUC cell injection. (D-J) IHC analysis of EGFP+ cells (D), CD4^+^(E), CD8^+^ (F), CD49b^+^ (G), Foxp3 (H), Ifng (I), and Cxcl9 (J) in breast tumors of female mice at 14^th^-days after ACT with SRC-3KO Tregs (SRC-3 KO) versus WT Tregs (Control). (K) Tumor luciferase activity in tumor-eradicated SRC-3^f/f^ female mice after ACT with SRC-3 KO Tregs and SRC-3^f/f^ female control mice after a 2^nd^ injection of E0771:LUC cells.

We next set up ACT cell dose-response experiments. In instances with a greater number of cells, ACT with SRC-3 KO Tregs (>600K cells) effectively eradicated tumors (Fig. 5B and Supplement Fig. S14C). However, ACT with lower doses of SRC-3 KO Tregs (2: 326K cells and 4: 230K cells) suppressed breast tumor growth compared to ACT with purified wild-type Tregs, but did not eradicate breast tumors (Fig. 5B and Supplement Fig. S14C) As expected, ACT with wild-type Tregs (800K cells) failed to block tumor growth (Fig. 5B and Supplement Fig. S14C). The repeat experiment with an increased number of animals (n=9) also confirmed that the low dose of SRC-3 KO Tregs (1: 230K cells, 3: 314K cells, 3: 358K cells) did not eradicate breast tumors compared to the high dose of SRC-3 KO Tregs (Fig. 5C). Therefore, the number (∼600K cells) of SRC-3 KO Tregs injected during ACT are critical factors for achieving complete breast tumor eradication in mice.

Breast tumor masses were still detected in E0771:LUC SRC-3^f/f^ recipient female mice up to 14 days after ACT with SRC-3 KO Tregs (Fig. 5A). To examine these tumors before their ultimate clearance, tumors were dissected from SRC-3 KO and WT ACT recipient animals 14 days after ACT cell injection. Because SRC-3 KO Tregs express EGFP, IHC with an EGFP antibody was used to trace the SRC-3 KO Tregs that infiltrated into the tumors of SRC-3^f/f^ host mice. As expected, EGFP expression was detected in the tumors of SRC-3^f/f^ female mice receiving ACT with SRC-3 KO Tregs, but not with wild-type Tregs (IHC control) (Fig. 5D and Supplement Fig. S15A). In contrast, IHC for effector immune cells revealed that ACT with SRC-3 KO Tregs significantly increased the number of CD4^+^ T cells, CD8^+^ T cells, and NK cells in cancers compared to ACT with wild-type Tregs (Figs. 5E-G and Supplement Fig. S15B-D). Additionally, ACT with SRC-3 KO Tregs markedly reduced Foxp3+ cells in breast tumors compared to ACT with WT Tregs (Fig. 5H and Supplement Fig. S15E). ACT with SRC-3 KO Tregs also significantly increased Ifng and Cxcl9 in breast tumors compared to ACT with wild-type Tregs (Fig. 5I and J and Supplement Figs. S15F and G). Our results obtained with ACT were in accordance with the observations from the SRC-3^d/d^:Treg female mice. Collectively, ACT with SRC-3 KO Tregs results in an increase in the infiltration of immune effector cells and enhances the Ifng/Cxcl9 axis in breast tumors, along with the subsequent generation of an efficient antitumor immune microenvironment that eradicated pre-existing E0771 tumors in SRC-3^f/f^ host mice.

Following tumor clearance to undetectable levels by ACT with SRC-3 KO Tregs, recurrence was not detected for at least 215 days (Fig. 5A). This observation suggested that ACT with SRC-3 KO Tregs leads to a state of long-term tumor resistance like that seen in SRC-3^d/d^:Treg female mice. To test this hypothesis, E0771:LUC cancer cells were ‘again reinjected’ into tumor-eradicated SRC-3^f/f^ female mice that had received prior ACT with SRC-3 KO Tregs (Supplement Fig. S13C). The reinjected E0771:LUC cells also did not develop into breast tumors in the tumor-eradicated SRC-3^f/f^ mice after ACT with SRC-3 KO Tregs, but did so in control SRC-3^f/f^ mice (Fig. 5K and Supplement Fig. S14D). Therefore, a single ACT injection with SRC-3 KO Tregs supports long-term tumor elimination and resistance in mice.

To define whether current immune checkpoint modulators have a similar tumor eradication effect as SRC-3 KO Tregs, E0771 breast tumor-bearing SRC-3^f/f^ female mice (C57/BL6J-Albino) were treated with an anti-PD-L1 antibody (7.5 mg/kg) or control IgG (7.5 mg/kg) twice a week for two weeks (Supplement Fig. S16A) based on a previous study (2). Compared to control IgG treatment, however, anti-PD-L1 antibody treatment did not suppress the growth of E0771 breast tumors in immune-intact mice (Supplement Figs. S16B and C). Therefore, anti-PD-L1 immunotherapy did not recapitulate the tumor eradication activity of SRC-3 KO Tregs.

## Discussion

Tregs play a critical role in cancer progression by generating an immune-suppressive microenvironment and thus have been a major focus for immune checkpoint inhibitor therapeutic development. While the vast majority of Treg-based immune checkpoint modulators are focused on cell membrane signaling proteins, some efforts to modulate Treg function at a transcriptional level within the nucleus have been pursued. For example, Treg-specific transcriptional coactivator p300 KO mice show an antitumor phenotype due to increased TCR-induced apoptosis of Tregs (28). However, Treg cell-specific p300 KO mice show incomplete tumor inhibition and develop adverse health effects, such as weight loss, dermatitis, lymphadenopathy, and splenomegaly related to Treg depletion as these animals age (28). In contrast, SRC-3 KO increases Treg proliferation and alters their function by switching them from protumor into antitumor immune cells while not causing systematic autoimmune disease, reproductive dysfunction, shortened lifespan, or lower body weight. These observations demonstrate a benign side-effect profile for SRC-3 KO Tregs in mice that contrasts with most immune checkpoint inhibitors and with Chimeric Antigen Receptor (CAR)-T cell therapies in that it avoids adverse effects such as cytokine release syndrome, neurological events, and other potentially serious side effects (29). Notably, our SRC-3-KO cells require only a ‘single injection’ to achieve a potent anti-tumor effect. Treg cell therapy, such as immunotherapy against co-inhibitory receptors (such as CTLA4, PD-L1, TIGIT, LAG3, Tim-3) and the targeting of co-stimulatory molecules (GITR, 4-1BB, OX40, CD27, and ICOS), effectively disrupt the immune suppressive function of Tregs and depletes Tregs cells to increase CTL function for tumor suppression (30). However, the systemic injection of Treg cell targeting antibodies leads to toxicity. For example, cancer immunotherapies such as CTLA-4 and PD-1 blockade are frequently accompanied by serious autoimmunity (31–33). However, our SRC-3 KO Treg therapy does not cause discernable toxicity. Even though spleen derived Foxp3^+^SRC-3^−^ Tregs cells possess a different gene expression profile than WT Tregs, spleen derived Foxp3^+^SRC-3^−^ Tregs did not cause any detectable phenotypic changes in SRC-3^d/d^:Treg female mice. Compared to that observed in the spleen however, tumors in SRC-3^d/d^:Treg female mice revealed new Foxp3^−^ SRC-3^−^ Tregs (25.6%). The elevated number of Foxp3^−^SRC-3^−^ Tregs in tumors was associated with increased Ifng levels within the tumor microenvironment. Thus, TCR-mediated engagement with tumor antigens likely induces the recruitment of Foxp3^−^SRC-3^−^ Tregs into tumors. The elevated Foxp3^−^SRC-3^−^ Tregs amplify an Ifng/Cxcl9 feedback loop in tumors that then actively recruit Cxcr3^+^ CTLs to actively generate a tumor-suppressive immune microenvironment that leads to cancer cell death. Therefore, tumor-specific generation of Foxp3^+^SRC-3^−^ Tregs effectively destroys tumors without generalized systemic cytotoxicity or autoimmunity. In future studies, we will investigate how TCR-mediated engagement with tumor antigens impacts Foxp3^−^SRC-3^−^ Treg function and what gene signature exists in Foxp3-SRC-3-Tregs compared to wild type Tregs.

SRC-3 KO Tregs express additional proteins including CRE-ERT2 recombinase and GFP proteins. In addition to this, E0771 breast tumors also express the luciferase protein. Although these non-mouse proteins possibly could act as neoantigens, and help the body mount an immune response (34, 35), these expressed exogenous genes have been successfully used in numerous cancer studies and appear not to be confounders in immune-related studies.

Another surprising but important therapeutic consideration related to the disruption of SRC-3 expression in Tregs is that ACT with SRC-3 KO Tregs eradicated tumors even though normal Tregs are present in SRC-3^f/f^ recipient mice. This observation raised the question of how SRC-3 KO Tregs take on a functionally dominant role over wild-type Tregs that leads to ACT-mediated tumor eradication. Our hypothesis is shown in Fig.6. The SRC-3 KO Tregs are more proliferative than WT Tregs in the spleen due to elevated levels of splenic TGF-β, Il-10, and IL35. Compared with WT Tregs, however, SRC-3 KO Tregs do not have heightened immune suppressive functions in the tumor microenvironment due to the down-regulation of many genes involved in immune suppression such as TIGIT and others. Furthermore, based on the homing activity of Tregs to cancer, such as from the CCL22/CCR4 axis (36, 37), SRC-3 KO Tregs can aggressively move into breast tumors. After TCR-mediated engagement with tumor antigens, SRC-3 KO Tregs, but not WT Tregs, produce high amounts of Ifng. The elevated levels of Ifng further induce antitumor immune activity in tumors, leading to increased CXCL9 expression from tumor cells; CXCL9 then recruits CXCR3+ immune cells, such as cytotoxic CD8+ T cells and NK cells, into cancers to increase effector immune cell infiltration (38, 39). Elevated Infg activates both recruited and resident CD8+ T cells and NK cells in breast tumors which can lead to the positive feedback that drives even more Infg production (23, 40). Furthermore, high Ifng production occurs only in the tumor microenvironment and is not elevated in the serum of the animals, avoiding systemic toxicity. Ifng is known to cause wild-type Treg cellular ‘fragility’ (41) along with high Treg production of perforin and granzymes (42). Consistent with this, our CD4 cell proliferation suppression assays indeed reveal that activated SRC-3 KO Tregs suppress WT Treg function. Therefore, the activated Ifng/Cxcl9 axis stimulated by SRC-3 KO Tregs would be expected to suppress WT Tregs in breast tumors by enhancing Ifng-induced cellular fragility in WT Tregs. Also, the upregulated Ifng/Cxcl9 axis can enhance the T-bet transcription factor in naïve CD4+ cells that supports their differentiation into T helper (Th)-1 cells while inhibiting their differentiation into Th-2 and Th17 cells (43, 44). The activated Ifng/Cxcl9 axis generated by SRC-3 KO Tregs also should stimulate the expansion of a proinflammatory Th-1-cell population within the tumor microenvironment to facilitate tumor eradication. Both the Treg cell-specific SRC-3 KO mouse and ACT with SRC-3 KO Tregs generated a strong tumor-resistance state. Breast tumor recurrence was never again detected, and subsequent 2^nd^ tumor injections did not induce cancers in Treg cell-specific SRC-3 KO mice and in mice receiving ACT with SRC-3 KO Tregs after eradication of the first tumor. Our data reveal that SRC-3 KO Tregs are more proliferative and viable than WT Tregs, and CD3/CD28 co-stimulation significantly increases the viability of SRC-3 KO Treg cells compared to WT Tregs. Thus, SRC-3 KO likely increases the life span of Tregs such that they can maintain tumor resistance in mice long-term. SRC-3 KO Tregs also showed increased expression of Il-10, Il-35, and Tgf-β compared to wild-type Tregs obtained from the spleen. TGF-β, IL-10, and IL-35 are known to stimulate the development of an induced regulatory T cell (iTreg) population (45–47). In addition, IL-10 enhances IL-2-stimulated proliferation of both CD4^+^ and CD8^+^ T cells by increasing cell division (48). Also, treatment with IL-35 has been shown to enhance the proliferation of Foxp3^+^CD39^+^ CD4^+^ T cells (49). Therefore, the elevation of Il-10, Il-35, and Tgf-β in SRC-3 KO Tregs is expected to increase/stabilize the number of SRC-3 KO Treg cells in the spleen by autocrine signaling pathways that then support their later infiltration into tumors, leading to tumor eradication.

**Fig. 6.**
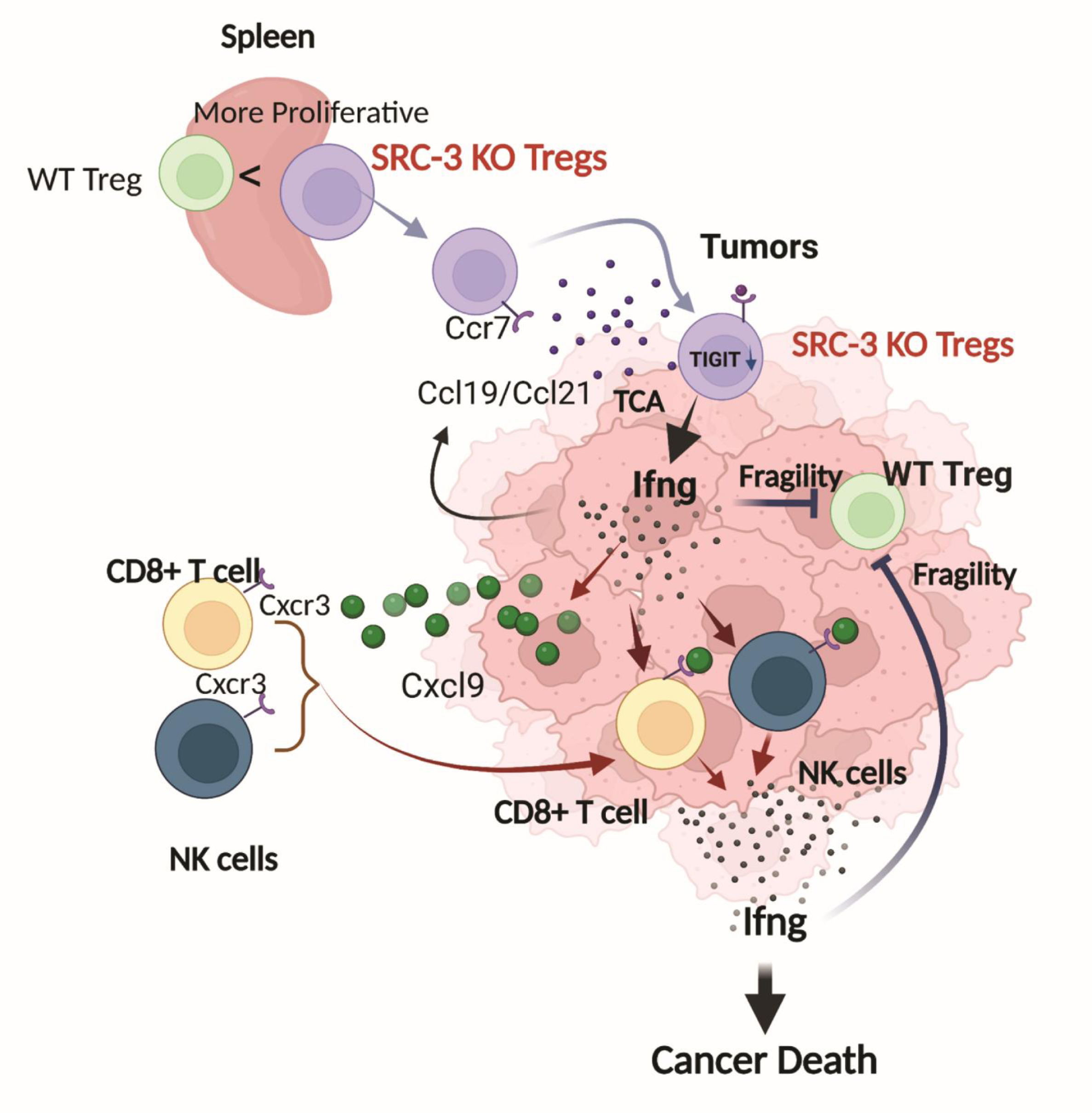
Model for tumor eradication by SRC-3 KO Tregs. SRC-3 KO Tregs are more proliferative than WT in spleens due to increased levels of Tgf-β, Il10, and IL35 in spleens. Compared to WT Tregs, however, SRC-3 KO Tregs have reduced immune suppressive function due to the down-regulation of genes involved in immune suppression, such as TIGIT. Based on the homing activity of Tregs to cancer (Ccl19/Ccl21/Ccr7 axis) (36, 37), SRC-3 KO Tregs more aggressively infiltrate into breast tumors. In breast tumors, SRC-3 KO Tregs induced Ifng expression based on TCR activation. The elevated Ifng causes the fragility of WT Tregs in breast tumors. Also, elevated Ifng induces Cxcl9 expression in breast tumors to actively recruit cytotoxic Cxcr3+ immune cells (such as CD8+ and NK cells) into breast tumors. In addition, the recruited and tumor resident CD8+ and NK cells activated by elevated Ifng eradicate cancer cells. The graphic was generated by BioRender.

As discussed in the introduction, SRC-3 is known to broadly function as a primary coactivator for numerous transcription factors in the cell (1). Given its known pleiotropic regulation of gene expression, it is not surprising to us that disruption of the SRC-3 gene results in dramatic changes in the Treg transcriptomic profile that affect the expression of key cytokines and membrane signaling molecules as a group. In addition, studies on the cell-autonomous roles of SRC-3 in cancer cells have revealed that this coactivator also broadly controls multiple transcriptional goal-oriented programs that drive key cancer-promoting pathways underlying cell proliferation, invasion, and metastasis (50–52). Because of these attributes of SRC-3 biology, further studies will need to consider the widespread transcriptional effects of this key coactivator on Treg function. Clearly, this pleiotropic effect cannot be expected to be restricted to the perturbation of only a single or even a few molecular pathways.

In conclusion, we demonstrate that SRC-3 KO Tregs show strong potential as a novel therapeutic approach for cancer therapy. SRC-3 KO Treg-based ACT therapy also avoids many of the known limitations seen for other immune-therapeutics, such as membrane immune checkpoint inhibitors, and is effective in breast/prostate mouse tumors where checkpoint inhibitor drugs are less effective. Also, it should be pointed out that current CAR-T therapy efficacy for solid tumor treatment has not yet been widely achieved, and significant side effects can limit their use (53). Since SRC-3 KO Tregs do not produce the strong, generalized autoimmune side effects that are typically seen with other immune checkpoint inhibitors, dose-dependent toxicities appear not to limit the dose of SRC-3 KO Tregs in ACT. Finally, SRC-3KO Tregs appear to confer extremely long-lasting protection against cancer recurrence in mice, suggesting that further work to develop genetically altered SRC-3 KO Tregs is warranted as a cell-based therapeutic agent to treat cancer.

## Materials and Methods

### Animal Studies

C57BL/6J (RRID:IMSR_JAX:000664) and Foxp3 ^tm9(EGFP/Cre/ERT2)Ayr^ mice (RRID:MGI:5288172), and Foxp3^tm4(YFP/icre)Ayr^/J (RRID:IMSR_JAX:016959) were purchased from Jackson Laboratory. Floxed SRC-3 (SRC-3^f/f^) mice employed to knock out the SRC-3 gene in specific tissues were described before (17). All animal studies were conducted with the approval of the Institutional Animal Care and Use Committee at Baylor College of Medicine. All mice were euthanized in a CO^2^ chamber. Additionally, any animals that showed a loss of mobility and weight loss were euthanized. Cancers were not allowed to grow to >20% of body weight or ulcerate. At the end of experiments, all animals were euthanized, and cancers were harvested at a volume of no more than 4,000 mm^3^.

### Animal numbers and power calculations

A tumor luciferase activity assay indicated that tumor luciferase activity was not detected in SRC-3^d/d^:Treg female mice compared to control mice (Figure 2A). Therefore, a power calculation based on these data revealed that the animal size (n=1/group) was sufficient to see a significant effect on tumor eradication by SRC-3 KO Treg cells in mice (p<0.001). We chose to use four or more mice in each group for all mouse experiments to achieve additional statistical power. Generation of Treg cell-specific SRC-3 KO mice. SRC-3^f/f^ mice originate from a mixed C57BL/6 x *129* genetic background (17). SRC-3^f/f^ female mice were crossed with Foxp3^tm4(YFP/icre)Ayr^/J male mice to generate Treg cell-specific SRC-3 KO bigenic mice (SRC-3^d/d^:Foxp3^tm4(YFP/icre)Ayr/+^). The genotyping of the SRC-3^f/f^ mice was conducted with primers (GAAACCTCAAGGTTATCCTTCAATT and TGCTCTGCTTAGATACCTGCTG) as previously reported (17). Genotyping of Foxp3^tm4(YFP/icre)Ayr/+^ mice was conducted using a protocol published by the Jackson Laboratory (https://www.jax.org/Protocol?stockNumber=016959&protocolID=36057). To generate the syngeneic mouse model of breast cancer with E0771 breast cells that spontaneously generated from C57BL/6J female mice, SRC-3^f/f^ female mice (mixed background) were mated with C57BL/6J male mice, and genotyping of the SRC-3^f/f^ mice were conducted with primers (GAAACCTCAAGGTTATCCTTCAATT and TGCTCTGCTTAGATACCTGCTG) as previously reported (17). After nine generations of mating to C57BL/6J males, SRC-3^f/f^ mice were established on a pure C57BL/6J background. Further, Jackson Laboratory confirmed that the genetic background of the Foxp3^tm9(EGFP/Cre/ERT2)Ayr^ (Foxp3^Cre-ERT2/+^) strain is in a pure C57BL/6J background. To further ensure a uniform genetic background, Foxp3^Cre-ERT2/+^ mice were also mated with C57BL/6J mice for nine generations to generate aFoxp3^Cre-ERT2/+^ mice in a pure C57BL/6J background. Genotyping of Foxp3^Cre-ERT2/+^ mice was conducted using a protocol published by the Jackson Laboratory (https://www.jax.org/Protocol?stockNumber=016961&protocolID=28173). SRC-3^f/f^ female mice were mated with Foxp3^Cre-ERT2/+^ male mice to produce SRC-3^f/f^:Foxp3^Cre-ERT2/+^ bigenic mice.

### Tamoxifen-Induced Treg SRC-3 gene KO in Treg cells

Tamoxifen (100 mg, Tocris Cat. No. 6432) was dissolved in 5 mL of corn oil (20 mg/mL). Then, 100 µl of the 20 mg/mL tamoxifen solution was injected intraperitoneally into SRC-3^f/f^:Foxp3^Cre-ERT2/+^ bigenic female mice (20g, eight weeks old) and female control mice (20g, eight weeks old), daily for five days. The final concentration of injected tamoxifen was 75 mg/kg.

### Determination of female reproductive toxicity resulting from Treg cell-specific SRC-3 KO

SRC-3^f/f^:Foxp3^Cre-ERT2/+^female mice (eight weeks old, n=5) and SRC-3^f/f^ female mice (eight weeks old, n=5) were treated with tamoxifen (75 mg/kg) daily for five days. Three weeks after tamoxifen treatment, each mouse in both groups was mated with a fertility-proven male mouse (C57BL/6J, eight weeks old) at a 1:1 ratio. The pregnancy rate and pup numbers per female mouse were then determined.

### Culture of the E0771 and RM-1cell lines

E0771 mouse mammary gland carcinoma cells (ATCC Cat# CRL-3461, RRID: CVCL_GR23) and RM-1 mouse prostate cancer cells (ATCC Cat# CRL-3310, RRID: CVCL_B459) cells were grown in RPMI 1640 containing 10% (vol/vol) fetal bovine serum (FBS) in a humidified incubator with 5% (vol/vol) CO^2^ at 37 °C.

### Generation of luciferase-labeled E0771 and RM-1 cells

The firefly luciferase cDNA was cloned into the PSMPUW-Hygro lentiviral vector (Cell Biolabs, catalog number: VPK-214). Lentiviruses containing the luciferase expression cassette were produced in 293 TN cells (System Bioscience, catalog number: LV900A-1) by transient transfection with the PSMPUW-Hygro lentiviral vector carrying the luciferase gene and the ViraSafe™ Lentiviral Packaging System (Cell Biolabs, catalog number: VPK-206) with Lipofectamine 2000 (Thermo Fisher Scientific, catalog number: 11668030). Recombinant lentivirus titer was measured using Lenti-XTM GoStixTM Plus (ClonTech, catalog number: 631280). E0771 and RM-1 cells were transduced with lentiviral vectors carrying the luciferase expression cassette with TransDux MAX^TM^ (System Bioscience, catalog number: LV860A-1). Luciferase-labeled E0771 and RM-1 cells were then selected in the presence of 300 µg/mL hygromycin. Luciferase gene expression in E0771 and RM-1 cells was validated using a luciferase activity assay kit (Promega). Luciferase-labeled E0771 and RM-1 cells were maintained in RPMI 1640 media supplemented with 10% FBS, penicillin/streptomycin, and 300 µg/mL hygromycin.

### Orthotopic injection of E0771:LUC cells into the mammary fat pads of syngeneic female recipient mice

A total of 1 × 10^5^ of E0771:LUC cells were resuspended in 50 µL of phosphate-buffered saline (PBS) and then injected into one of the 4^th^ mammary fat pads in SRC-3^f/f^ (eight weeks old, n=8) and SRC-3^d/d^:Treg (eight weeks old, n=8), as described in our previous studies (54). Tumor growth was monitored by evaluating luciferase activity using an *In Vivo* Imaging System (IVIS, Xenogen). Luciferase activity in E0771 tumors was assessed twice a week using IVIS.

### Orthotopic injection of luciferase-labeled prostate cancer cells into the prostates of syngeneic male recipient mice

A total of 1 × 10^3^ luciferase-labeled RM-1(RM-1:LUC) cells were resuspended in 50 µL of phosphate-buffered saline (PBS) and then orthotopically injected into the ventral lobe of the prostate in SRC-3^f/f^ (n=3, eight weeks old) and SRC-3^d/d^:Treg male mice (n=3, eight weeks old), as described in previous studies (55). Tumor growth was monitored by evaluating luciferase activity using an IVIS imaging system. Luciferase activity in RM-1 prostate cancers was assessed twice a week.

### IVIS analysis of luciferase bioluminescence in mice

Mice were anesthetized with a 1.5% isoflurane/air mixture using an inhalation anesthesia system (VetEquip). Next, D-Luciferin (Xenogen, Cat# 122799) was injected intraperitoneally at 40 mg/kg mouse body weight. Ten minutes after D-luciferin injection, mice were imaged using IVIS under continuous exposure of 1% to 2% isoflurane. Imaging parameters were maintained for comparative analysis. Grayscale and pseudocolor images showing bioluminescence were superimposed and analyzed using Living Image software (Version 4.4, Xenogen). A region of interest (ROI) was manually selected over the relevant areas showing a luciferase signal. The ROI area was kept constant across the experiments, and the luciferase activity was recorded as total luciferase photon counts per second per cm^2^ within the ROI.

### Prevention of tumor initiation in SRC-3^d/d^:Treg female mice

SRC-3^f/f^:Foxp3^Cre-ERT2/+^ (eight weeks old,n=2) and SRC-3^f/f^ female mice (eight weeks old, n=2) were treated with tamoxifen (75 mg/kg) daily for five days. After two weeks, E0771:LUC (1×10^5^) cells were orthotopically injected into the mammary fad fat of one SRC-3^d/d^:Treg female mouse. As controls, E0771:LUC (1×10^5^ cells) were orthotopically injected into one SRC-3^f/f^ female mice. Tumor luciferase activity was determined twice a week by IVIS imaging.

### Repression of pre-existing E0771 tumors in mice

SRC-3^f/f^:Foxp3^Cre-ERT2/+^ (eight weeks old, n=3) and SRC-3^f/f^ female mice (eight weeks old, n=4) were orthotopically injected with 1×10^5^ E0771:LUC cells per mouse. On day seven, after cell injections, tumor-bearing SRC-3^f/f^:Foxp3^Cre-ERT2/+^ and SRC-3^f/f^ female mice were treated with tamoxifen (75 mg/kg) daily for five days. Afterward, the tumor luciferase activity was determined twice a week by IVIS imaging.

### Tamoxifen and vehicle treatment of E0771 breast tumor-bearing SRC-3^f/f^ female mice

E0771 breast cancer cells (1×10^5^ cells) were orthotopically injected into SRC-3^f/f^ female mice (8 weeks old). On the 9^th^ day after breast cancer cell injection, tumor-bearing mice were treated with tamoxifen (75 mg/kg, once a day for 5 days) or vehicle as controls. Tumor luciferase activity was determined with an IVIS bioluminescent imager and quantified with Living Image software (version 4.7.4).

### Injection of a second round of E0771:LUC cells into tumor-eradicated SRC-3^d/d^:Treg female mice

Seven weeks after the first injection of E0771:LUC, tumor-eradicated SRC-3^d/d^:Treg female mice (Figure 2A, n=5) were orthotopically injected with E0771:LUC cells (1 x10^5^) in their mammary fat pads. As controls, SRC-3^f/f^ female mice (eight weeks old, n=5) were orthotopically injected with E0771:LUC cells (1 x10^5^) into their mammary fat pads. Luciferase activity of tumors was determined twice a week by IVIS imaging.

### ACT of Treg cells

The SRC-3^f/f^:Foxp3^Cre-ERT2/+^ bigenic female mice (eight weeks old, n=9) and SRC-3^f/f^ female mice (eight weeks old, n=9) were treated with tamoxifen (75 mg/kg) daily for five days. After two weeks, SRC-3 KO Tregs were isolated from the spleens of SRC-3^d/d^:Treg mice. As controls, wild-type Tregs were isolated from the spleens of SRC-3^f/f^ mice treated with tamoxifen. The SRC-3^f/f^ mice (eight weeks old, n=18) were orthotopically injected with E0771:LUC cells (1 x10^5^) into mammary fat pads. Seven days after injection of E0771:LUC cells, isolated SRC-3 KO Tregs (cell number: 200 ∼ 800 K cells) in 50 µL of MDEM/F12 media were given to tumor-bearing littermate SRC-3^f/f^ mice by retro-orbital injection (n=9). As controls, wild-type Tregs (cell number: ∼900 K cells, cell viability: ∼90%) in 50 µL of MDEM/F12 media were given to tumor-bearing littermate SRC-3^f/f^ female mice by retro-orbital injection (n=9). After ACT, tumor luciferase activity was determined in mice adoptively transferred with SRC-3 KO versus wild-type Tregs.

### Second round of E0771 cell injections into SRC-3 KO ACT Treg recipient mice

Forty days after the first round of E0771 cell injections, tumor-eradicated SRC-3^f/f^ mice that received ACT of SRC-3 KO Tregs (n=3) were orthotopically injected with E0771:LUC cells (1×10^5^) into a mammary fat pad. As controls, SRC-3^f/f^ female mice (eight weeks old, n=2) were orthotopically injected with E0771 cells (1×10^5^) into a mammary fat pad. Tumor luciferase activity was determined twice a week by IVIS imaging.

### IHC analysis

Fresh mouse tumor tissues were fixed with 10% buffered formalin and then embedded in paraffin for subsequent sectioning. Tissue slices were deparaffinized by sequential soaking in xylene, ethanol, and water. Antigen retrieval was performed with antigen unmasking solution (Vector Laboratory, catalog number: H-3300). Endogenous peroxidase activity was blocked with a 3% (vol/vol) hydrogen peroxide solution. Sections were blocked with 2.5% (w/v) normal goat serum to reduce nonspecific antibody binding. Slices were incubated with primary antibodies, including anti-CD4 (Abcam Cat# ab183685, RRID:AB_2686917, 1:300 dilution), and anti-CD8 (Abcam Cat# ab217344, RRID:AB_2890649, 1:300 dilution), anti-CD49b (Abcam Cat# ab181548, RRID:AB_2847852, 1:300 dilution), IFNG (rabbit, Novus, catalog number: NBP2-66900, 1:100 dilution), CXCL9 (Thermo Fisher Scientific Cat# PA5-81371, RRID:AB_2788585, 1:100 dilution), EGFP (Bioss Cat# bs-2194R, RRID:AB_10881247, 1:100 dilution), and anti-Foxp3 (Thermo Fisher Scientific Cat# PA1-16876, RRID:AB_568540, 1:300 dilution) antibodies at 4 °C overnight. Slices were then stained with HRP polymer-conjugated secondary antibodies (Vector Laboratories, catalog number: 7401 for rabbit antibodies). Slices were reacted with freshly prepared DAB solution (Dako, catalog number: K3468). After staining, nuclear counterstaining was performed with Mayer’s hematoxylin. Quantification of IHC signal was conducted with QuPath software (56).

### Dual Immunofluorescence Staining

Tissue slices were deparaffinized by sequential soaking in xylene, ethanol and water. Antigen retrieval was performed with antigen unmasking solution (Vector Laboratory, catalog number: H-3300). Slides were blocked with 2.5% (w/v) normal goat serum to reduce nonspecific antibody binding. Slices were incubated with anti-SRC-3 (Cell Signaling Technology Cat# 2126, RRID:AB_823642, 1:100 dilution) and anti-FOXP3 (rat, LSBio, catalog number: LS-C344878, 1:100 dilution) antibodies at 4 °C overnight. Afterward, slides were washed with TBST [20 mM Tris (pH 7.5), 150 mM NaCl and 0.1% (w/v) Tween 20] and then incubated with goat anti-rabbit Alexa Fluor 488 (Thermo Fisher Scientific Cat# A-11008, RRID:AB_143165, 1:500 dilution) and goat anti-rat Alexa Fluor 594 (Thermo Fisher Scientific Cat# A-11007, RRID:AB_10561522, 1:500 dilution) for 1 h at room temperature. Afterward, slices were washed with TBST and mounted with Antifade Mounting Media containing DAPI (Vector, catalog number: H-2000).

### Mouse cytokine/chemokine analysis with tumors

Breast tumors were isolated from SRC-3^f/f^ (n=4) and SRC-3^d/d^:Treg female mice (n=4) on day 14 after E0771 cell injection. Cytokine and chemokine levels in breast cancers were determined using Proteomic Profile Mouse Cytokine Array Kit (R&D System, ARY0066). Afterward, cytokine levels were quantified with the ImageJ program (57).

### Mouse splenocytes isolation and immunophenotyping

Single mouse spleens were suspended in 2.6 mL of RPMI1640 media (10% FBS, 1% anti/anti) in C-tubes. Each tube was treated with 375 µg of DNAase I (ThermoFisher, Cat# EN0521) and 1.87mg of Collagenase IV (Sigma-Aldrich, Cat# C4-28) in 300 µL of total media and placed on GentleMACS dissociator (Miltenyi Biotec, Cat# 130-095-235) for 15 mins (2x after 10 mins interval). Then, 400 µL of stopping buffer (1X PBS, 0.1M EDTA) was added and centrifuged for a minute at 100 rpm to collect the liquid at the bottom. The resulting single cell suspension was filtered (40 µm mesh), RBC lysed using 200 µL of 1X Ebioscience Lysis solution (Thermo fisher catalog number: 00-4300-54) for five minutes and centrifuged for six minutes at 600x g. The supernatant was discarded, and the cell pellet was suspended in 100 µL of complete media, and 50 µL of single cell suspension was used for cell counting using Vi-cell. Each sample was Fc blocked before staining using 50 µL of 2X FC Block Solution (FACS Buffer, 1:125 dilution of anti-CD16/CD32 (BD) and incubated for 10 minutes on ice, 100 µL of cold FACS buffer was added and centrifuged for six mins at 500x g. The pellet was suspended in FACS buffer and stained using appropriate antibodies for panel 1 (lymphoid panel) and panel 2 (myeloid panel). Single-cell splenocyte suspensions were analyzed using a BD LSRII for the antibody panel by the Cell Cytometry and Sorting Core at Baylor College of Medicine in collaboration with the International Mouse Phenotyping Consortium. Surface antibodies included in this study are—CD5, CD4, CD44, CD8, CD25, CD161, CD62L, CD19, Ly6C, Ly6G, CD21, CD35, CD11b, CD11c, IA-E and CD23.

### Mouse splenocyte and Treg cell isolation

Each mouse spleen was placed over a 70 µm filter on top of 50 mL tubes. Mouse spleens were processed through the filter using a syringe with RPMI1640 supplemented with 10% FBS. The resultant cell suspension was incubated with 1x Ebioscience RBC lysis buffer for five minutes on ice and then filtered through a 40 µm mesh to obtain single-cell splenocytes. Splenocytes were counted using a Vi-cell counter. Finally, T cell isolation was performed using Miltenyi Biotec’s MACS CD4^+^CD25^+^ T cell isolation kit using the manufacturer’s recommended protocol. In this process, positive selection of CD4 ^+^ CD25 ^+^ (Tregs) cells was performed using MACS CD25 MicroBeads and non-selected cells were labeled as CD4 ^+^ CD25^−^ (Tconv) cells.

### Flow Cytometry Analysis for SRC-3 KO Tregs

Spleens and tumors were dissected from mice and rinsed with cold PBS. Spleens were directly mashed through a 100 μm filter using the plunger from a 10 ml syringe. After centrifuge, the pellet was incubated with RBC lysis buffer for 5 min on ice to remove red blood cells (RBC) and then used for Treg isolation with a CD4+CD25+ Treg isolation kit (Miltenyi Biotec, cat # 130-091-041), according to the manufacturer’s instructions. Isolated Tregs were fixed and permeabilized using the Foxp3 Transcription Factor Fixation/Permeabilization buffer (eBioscience) and then incubated with unconjugated anti-FOXP3 (Cell Signaling Inc.) antibody and anti-SRC-3-Alexa647 (sc-5305, Santa Cruz Biotech) at 4°C overnight. The next day, Foxp3 protein was labeled with a fluorescent secondary antibody. Cells were washed twice with PBS before flow analysis. Tumor tissue was first minced with scissors and then digested with collagenase IV (3 mg/ml) in RPMI 1640 medium containing 10% FBS at 37°C for one hour. Digestion was then further performed by mashing and then filtered through a 40 μm filter using a 10 ml syringe plunger to get a single-cell suspension. After lysis of RBCs, remaining cells were processed for T cell purification using the Pan T cell isolation kit (<u>Miltenyi Biotec</u>, cat # 130-095-130). Cell viability of the purified T cells was ascertained by labeling with the eBioscience™ Fixable Viability Dye eFluor™ 506 for 30 min on ice. T cells were first processed for surface marker labeling by incubation with anti-CD8-BV421 and anti-CD4-BV650 (Biolegend) for 30 min on ice. To stain the nuclear proteins Foxp3 and SRC-3, cells were first fixed and permeabilized using the Foxp3 Transcription Factor Fixation/Permeabilization buffer (eBioscience) and then incubated with unconjugated anti-FOXP3 and the fluor conjugated anti-SRC-3-Alexa647 antibodies at 4°C overnight. The next day, Foxp3 protein was labeled with a fluorescent secondary antibody. Cells were washed twice with PBS before flow analysis. Flow cytometry acquisition was performed on an LSRII cytometer (BD), and the collected data was analyzed using FlowJo v10.8.1 software (TreeStar).

### RNAseq analysis of SRC-3 KO Tregs

Tregs were isolated from spleens from SRC-3^f/f^ female mice and Treg cell-specific SRC-3 KO bigenic mice (SRC-3^d/d^:Foxp3^tm4(YFP/icre)Ayr/+^). RNA was isolated from Treg cells using a Qiagen RNAasy micro-RNA isolation kit. The isolated RNA was tested for quality on an Agilent 2100 Bioanalyzer and quantitated using a NanoDrop Spectrophotometer. Low-input RNA-seq was conducted using a Takara SMART-seq v4 ultra-low-input RNA kit using the manufacturer’s recommended protocol. In brief, purified RNA was incubated with lysis buffer for five minutes, 3-SMART-seq CDS primer II and V4 oligonucleotides were added for first stranded cDNA synthesis. cDNA was amplified using PCR Primer II A and subsequently purified using Ampure XP beads (Beckman). Illumina libraries were prepared using Nextera XT DNA library preparation kits (Illumina) and sequenced using an Illumina Novaseq 6000. The RNA-seq data is pair-end sequenced with 150 bp read depth. The raw FASTAseq files were imported to the NGS biodata base. The cellular pathways enriched in SRC-3 KO Treg cells compared to WT Treg cells were analyzed with the Reactome pathway database.

### RNA expression analysis of genes of interest in SRC-3 KO versus WT Treg cells

Total mRNA was isolated from Tregs from the spleens and breast tumors of breast tumor-bearing SRC-3^f/f^ and SRC-3^d/d^:Treg female mice sing TRI reagent (Sigma Aldrich T9424) and the GlycoBlue coprecipitation protocol. cDNA was generated from total Treg RNA (∼0.5 to 10 μg) using a Vilo SuperScript Master Mix cDNA synthesis kit (Invitrogen catalog number: 11754050). To study relative gene expression, qPCR was carried out using sequence-specific primers and with PowerTrack SYBR green master mix (ThermoFisher, catalog number: A46109). Relative mRNA expression was calculated by the 2^−ΔΔCT^ method of quantitative PCR after normalization with 18S rRNA levels. At least three technical replicates represent each result. Primer sequences are as follows for each cytokine: SRC-3 exon 11 (GTCCCAACCAGCAGAACATC, GAAGCAAAGGAAAACGCAGC), Il-2ra (GGATGGGAATCACAAAGCTC, CCAGGGATCAGAAGGAAACA), Ifng (CGGCACAGTCATTGAAAGCCTA, GTTGCTGATGGCCTGATTGTC), Il-10 (AAGGCAGTGGAGCAGGTGAA, CCAGCAGACTCAATACACAC), Il-35 (GCTCCCCTGGTTACACTGAA, ACGGGATACCGAGAAGCAT), Tgfβ (TGATACGCCTGAGTGGCTGTCT, CACAAGAGCAGTGAGCGCTGAA), Tigit (GAATGGAACCTGAGGAGTCTCT, AGCAATGAAGCTCTCTAGGCT), Klrbic (ATGGACACAGCAAGTATCTACCT, AGCTCTCAGGAGTCACTTTATCT), Klrk1 (ACTCAGAGATGAGCAAATGCCATAA, CAGGTTGACTGGTAGTTAGTGCTAAT), Ccr7 (GACCCAGGTGTGCTTCTG, GGCCCAGAAGGGAAGAATTAG), and 18S rRNA (TCCGATAACGAACGAGACTC, CAGGGACTTAATCAACGCAA)

### RT-PCR for Treg chemotactic ligands in E0771 cells upon Ifng treatment

E0771 breast cancer cells were cultured with RPMI1640 media containing 10% FBS. When cell confluence was 90%, E0771 breast cells were treated with 0, 1, 10, and 100 ng/ml of active mouse recombinant Ifng protein (Abcam, catalog number: 259378) for 3 days. RPMI1640 media containing Ifng were exchanged every other day. Total mRNA was isolated from E0771 breast cancer cells treated with Ifng using a RNeasy Kit (Qiagen, catalog number: 74104). The cDNA was generated from total Treg RNA (∼0.5 to 10 μg) using a Vilo SuperScript Master Mix cDNA synthesis kit (Invitrogen, catalog number: 11754050). To study relative gene expression, qPCR was carried out using sequence-specific primers and PowerTrack SYBR green master mix (ThermoFisher, catalog number: A46109). Relative mRNA expression was calculated by the 2^−ΔΔCT^ method of quantitative PCR after normalization with 18S rRNA levels. At least three technical replicates represent each result. Primer sequences are as follows for each cytokine: Ccl2 (GCTCAGCCAGATGCAGTTA, TCACACTGGTCACTCCTACA), Ccl20 (TTGCTTTGGCATGGGTACT, ACTCTTAGGCTGAGGAGGTT), Ccl19 (GGTGCTAATGATGCGGAAGA, GCTGTTGCCTTTGTTCTTGG), Ccl21 (TGCAAGAGAACTGAACAGACAC, TTCCCTGGGAGACACTCTTT), Ccl11 (ACCTTGTGCAGGCAGTTT, GGATGGAGCCTGGGTGA), and 18S rRNA(TCCGATAACGAACGAGACTC, CAGGGACTTAATCAACGCAA).

### Determination of Cxcl9 expression in E0771 cells upon Ifng treatment

E0771 breast cancer cells were cultured with RPMI1640 media containing 10% FBS. When cell confluence was 90%, E0771 breast cells were treated with 0 and 100 ng/ml of active mouse recombinant Ifng protein (Abcam, catalog number: 259378) for 24, 48, and 72 hours. RPMI1640 media containing Ifng were exchanged every other day. As described above, total mRNA and cDNA were generated from E0771 breast cancer cells treated with Ifng. The RNA level of Cxcl9 was determined with primer set (ACTCCAACACAGTGACTCAATAG, CGTTCTTCAGTGTAGCAATGATTT) compared to 18S rRNA (TCCGATAACGAACGAGACTC, CAGGGACTTAATCAACGCAA).

### Anti-Infg antibody treatment of breast tumor-bearing SRC-3^f/f^ and SRC-3^d/d^ :Foxp3^Cre/+^ female mice

SRC-3^f/f^ and SRC-3^f/f^:Foxp3^Cre/+^ female mice (8 weeks old) were treated with tamoxifen (75 mg/kg, once a day for 5 days). At the end of the 2^nd^ week after the final tamoxifen treatment, SRC-3^f/f^ and SRC-3^d/d^:Treg female mice were intraperitoneally treated with an anti-mouse Ifng antibody (BioXCell, catalog number: DE0055, 0.2 mg/kg, twice a week for 2 weeks) or rat IgG (BioXCell, catalog number: BE00884, 0.2 mg/kg, twice a week for 2 weeks) as controls. On the third day after the first antibody treatment, E0771 cells (1x 10^5^ cells) were orthotopically injected into the mammary glands of SRC-3^f/f^ and SRC-3^d/d^:Treg female mice. Tumor luciferase activity was determined with a IVIS bioluminescent imager.

### Anti-PD-L1 antibody treatment of E0771 breast tumor-bearing SRC-3^f/f^ female mice

E0771 breast cancer cells (1×10^5^ cells) were orthotopically injected into SRC-3^f/f^ female mice (8 weeks old). On the ninth day after breast cancer cell injection, tumor-bearing mice were treated with anti-PD-L1 antibody (100 μg/20g mouse, every third day totaling 4 treatments) and control rat IgG (100 μg/20g mouse, every third day totaling 4 treatments) based on a previous study (58). Tumor luciferase activity was determined with an IVIS bioluminescent imager. The luciferase activity was quantified with Living Image (version 4.7.4) software.

### Treg Proliferation assay

Treg cells were purified from splenocytes collected from the SRC-3^f/f^:Foxp3^Cre-ERT2/+^ bigenic and SRC-3^f/f^ mice two weeks after the last injection of tamoxifen, using a Miltenyi Biotec’s MACS CD4+CD25+ T cell isolation kit according to the manufacturer’s protocol. Tregs were seeded to anti-CD3 antibody pre-coated 48-well plates at a density of 3×10^5^cells/ml in the stimulation medium (RPMI+10%FBS+1%anti-anti and supplemented with 2µg/ml anti-CD28 antibody and 1000 U/ml mouse IL-2) for two days. On day 3, half of the medium was replaced with fresh maintenance medium (RPMI+10%FBS+1% anti-CD28 and supplemented with 1000 U/ml mouse IL-2). Cell viability and density was detected by trypan blue staining on days 0, 3, 5 and 7 with the Bio-Rad TC20 cell counter.

### CD4 cell proliferation suppression assays (mixed Treg and Tconv experiments)

Tregs and Tconv cells were isolated as described in the “Mouse splenocytes and Treg cell isolation” section. CD4+CD25-(Tconv) cells were washed with 1X PBS and then incubated with Cell Trace Violet (CTV, Thermo-Fisher, catalog # C34557) solution in 1X PBS at a concentration of 5 mM per 1×10^6^ cells for 30 minutes at 37°C for staining. Serum-enriched media was added to quench any unreacted CTV and then centrifuged at 300 rpm for five minutes at 40 °C. Stained Tconv cells were resuspended in enriched activation media (RPMI1640 + 10% FBS + 50 μM mercapto ethanol + 2 μM CD28 (0.5 mg/mL) + 200 U IL2 (100 μg/mL) at a concentration of 5×10^4^ cells/mL. Unstained Tregs were resuspended in enriched activation media at a concentration of 1×10^6^ cells/mL. Tregs were mixed with stained Tconv cells at different ratios and then plated in a CD3 (5 μg/mL) precoated U-bottom 96 well-bottom plate with incubation at 37°C. The proliferation of stained Tconv cells was measured using flow cytometry on days 3, 4, and 5.

### Quantification and statistical analysis

Data are presented as the mean ± SEM values. Statistical comparisons between two groups were performed with one-way ANOVA, followed by the Student’s t-test or the Mann– Whitney U-test. Sidak’s multiple comparison test was used to compare more than two groups. A p-value < 0.05 was considered significant. Prism 8.0 software was used for statistical analyses. The differential RNA expression profile between SRC-3 KO Treg versus wild-type Treg was analyzed using R, Bioconductor, and orange program. The cellular pathways associated with SRC-3 KO Treg cells were analyzed with DAVID and GSEA programs.

## Acknowledgments

We gratefully thank Drs. Sandra L. Grimm and Christian Coarfa for the assistance with RNA-seq data analysis. This work is partly supported by funding from the National Institutes of Health grants R01 HD07857 and R01 HD08188 (BWO), and R01HD098059 (SJH).

## Abbreviations

SRC-3 Steroid Receptor coactivator-3

ACT: Adoptive cell transfer
Ccl: Chemokine (C-C motif) ligand
Ccr: Chemokine (C-C motif) Receptor
CTL: Cytotoxic T lymphocytes
Cxcl: C-X-C Motif Chemokine Ligand
CXCR: C-X-C chemokine receptor
EGFP: Enhanced green fluorescent protein
Foxp3: Forkhead box transcription factor 3
H&E: Hematoxylin and eosin
Ifng: Interferon-γ IL Interleukin
Il-1ra: Interleukin 1 receptor antagonist
IVIS: In Vivo Image System
Klrb1c: Killer cell lectin-like receptor subfamily B, member 1,
Klrk1: Killer Cell Lectin Like Receptor K1
KO: Knockout LUC Luciferase
Mip: Macrophage Inflammatory Proteins NS. Nonspecific
NK.: Natural Killer
PLC: Percentage of Labelled Cells TCR T Cell Receptor
TGF-β: Transforming growth factor beta
Tigit: T Cell Immunoreceptor With Ig And ITIM Domains
Timp-1: Tissue inhibitor matrix metalloproteinase 1
Treg: Regulatory T cells

## Data and code availability

The accession numbers for the RNA-seq data from SRC-3 KO and wild-type Tregs in this paper are GEO: GSE216931.

## Supplement Figure Legends

**Supplement Fig. S1.**
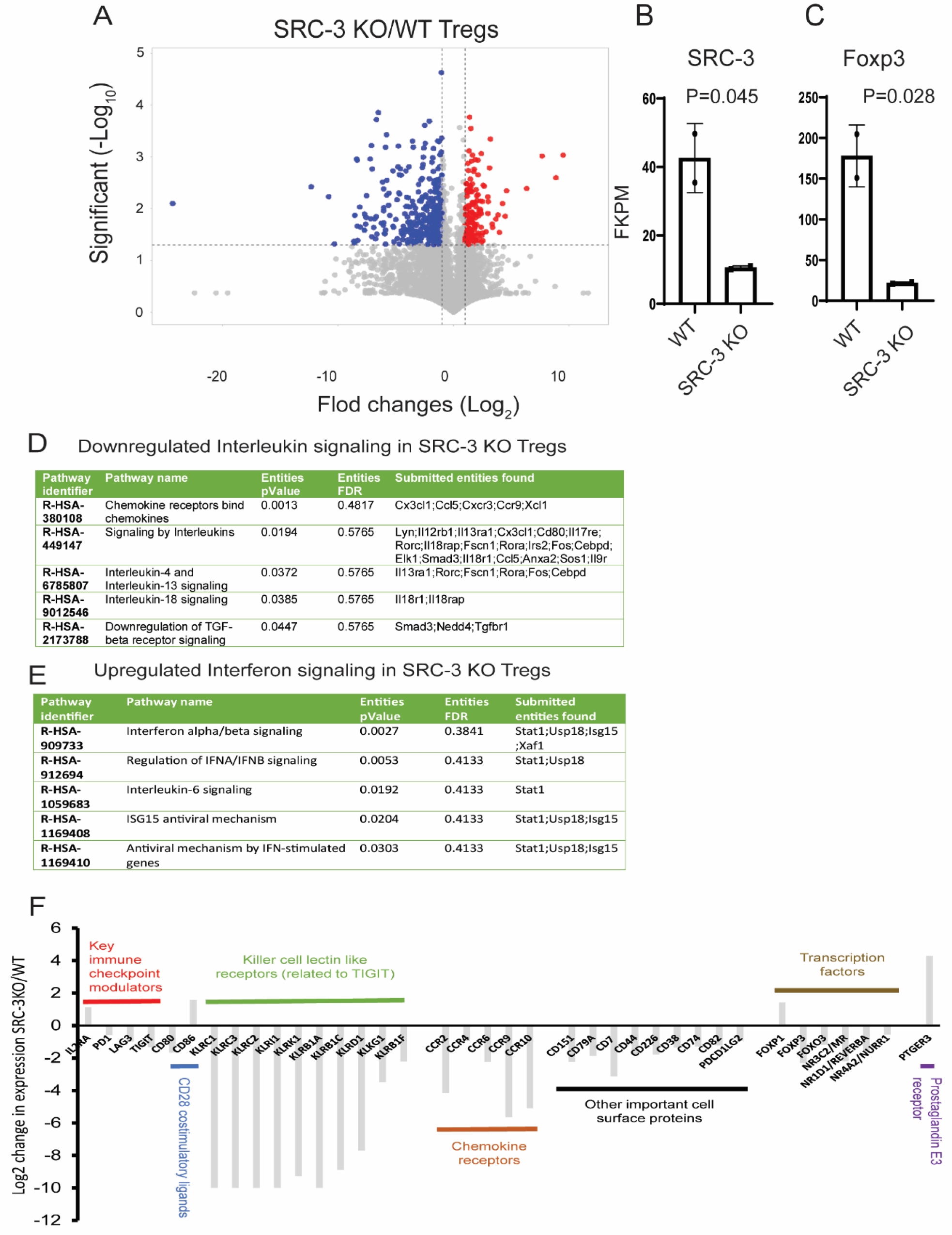
Downregulation of immune genes in SRC-3 KO Tregs. (A) Volcano plot analysis of differential RNA expression between SRC-3 KO Tregs from SRC-3^d/d^:Foxp3^tm4(YFP/icre)Ayr/+^ female mice and WT Tregs from SRC-3^f/f^ female mice determined by RNA-seq analysis. Genes colored in red were elevated in SRC-3 KO Tregs (> 2-fold, P<0.05), and genes colored with blue were significantly down-regulated in SRC-3 KO Tregs (> 2-fold, P<0.05) compared to WT Tregs. (B and C) SRC-3 (B) and Foxp3 (C) levels in between WT versus SRC-3 KO Tregs. (D) Downregulated interleukin signaling in SRC-3 KO Tregs. (E) Upregulated Interferon signaling in SRC-3 KO Tregs. (F) Dysregulated gene expression profile of SRC-3 KO Tregs compared to WT Tregs. The relative fold change of each target gene was calculated from the ratio of SRC-3 KO Treg to WT Treg mRNA expression.

**Supplement Fig. S2.**
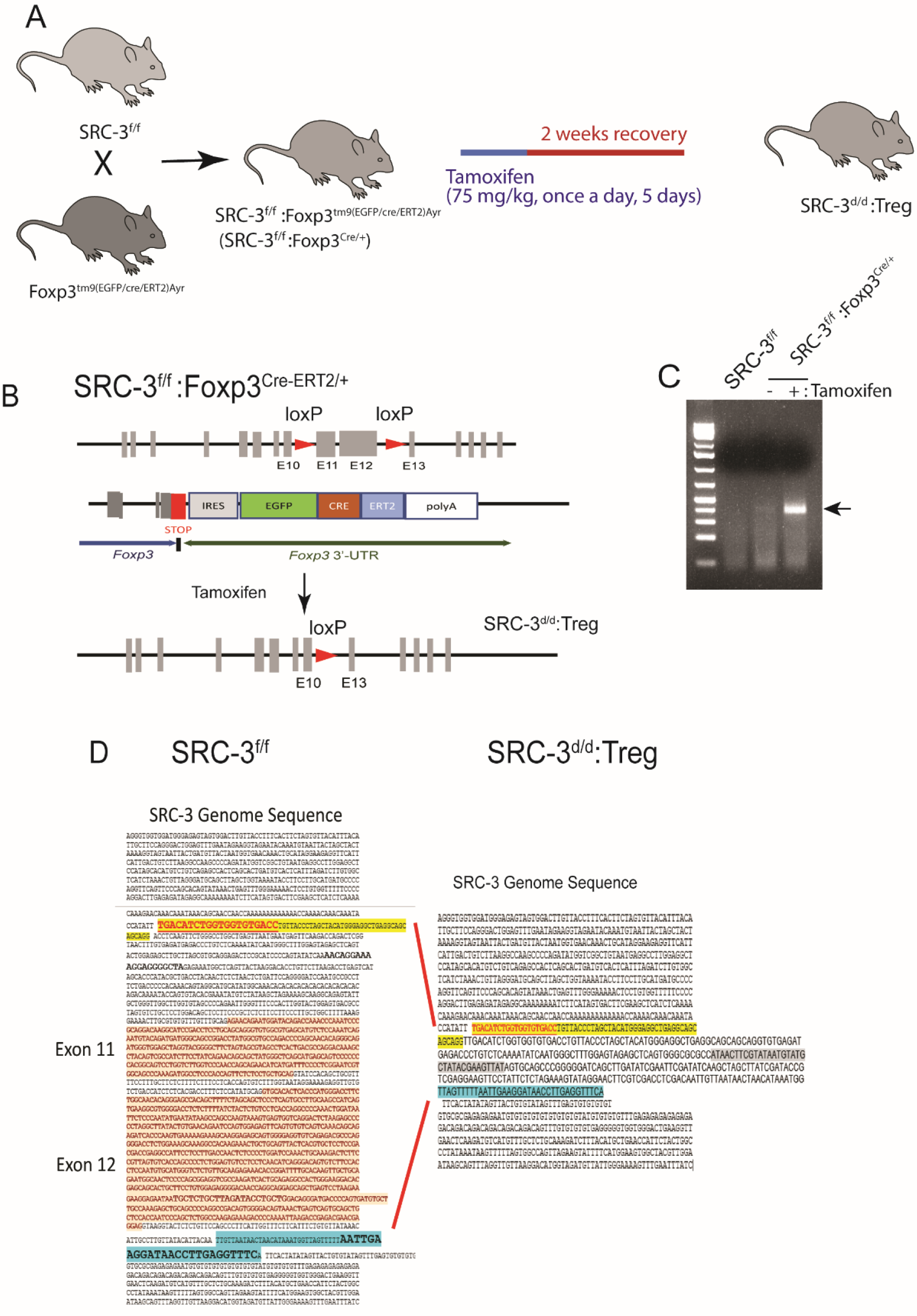
Generation of Treg cell-specific SRC-3 KO mice. (A) Schematic diagram of the generation of Treg cell-specific SRC-3 KO mice (SRC-3^d/d^:Treg). (B) Diagram of SRC-3 gene deletion in Tregs. Cre-ERT2 deletes exons 11 and 12 in the SRC-3^f/f^ mouse specifically in Foxp3 expressing cells upon tamoxifen treatment under the control of the Foxp3^tm9(EGFP/Cre/ERT2)Ayr^ allele in bigenic mice. (C) Genotyping PCR with genomic DNA from Tregs obtained from the spleen of SRC-3^f/f^ and SRC-3^f/f^:Foxp3^Cre-ERT2/+^ mice treated with either tamoxifen or vehicle and with primers mapping to exon 11 of the SRC-3 gene. (D) DNA sequence analysis of exon 11 and exon 12 showing deletion of these exons in the SRC-3 gene. The PCR product in Panel (C) was isolated and sequenced. Exon 11 and exon 12 (orange highlight) were deleted, and one loxP site (highlighted in gray) exists in the PCR product.

**Supplement Fig. S3.**
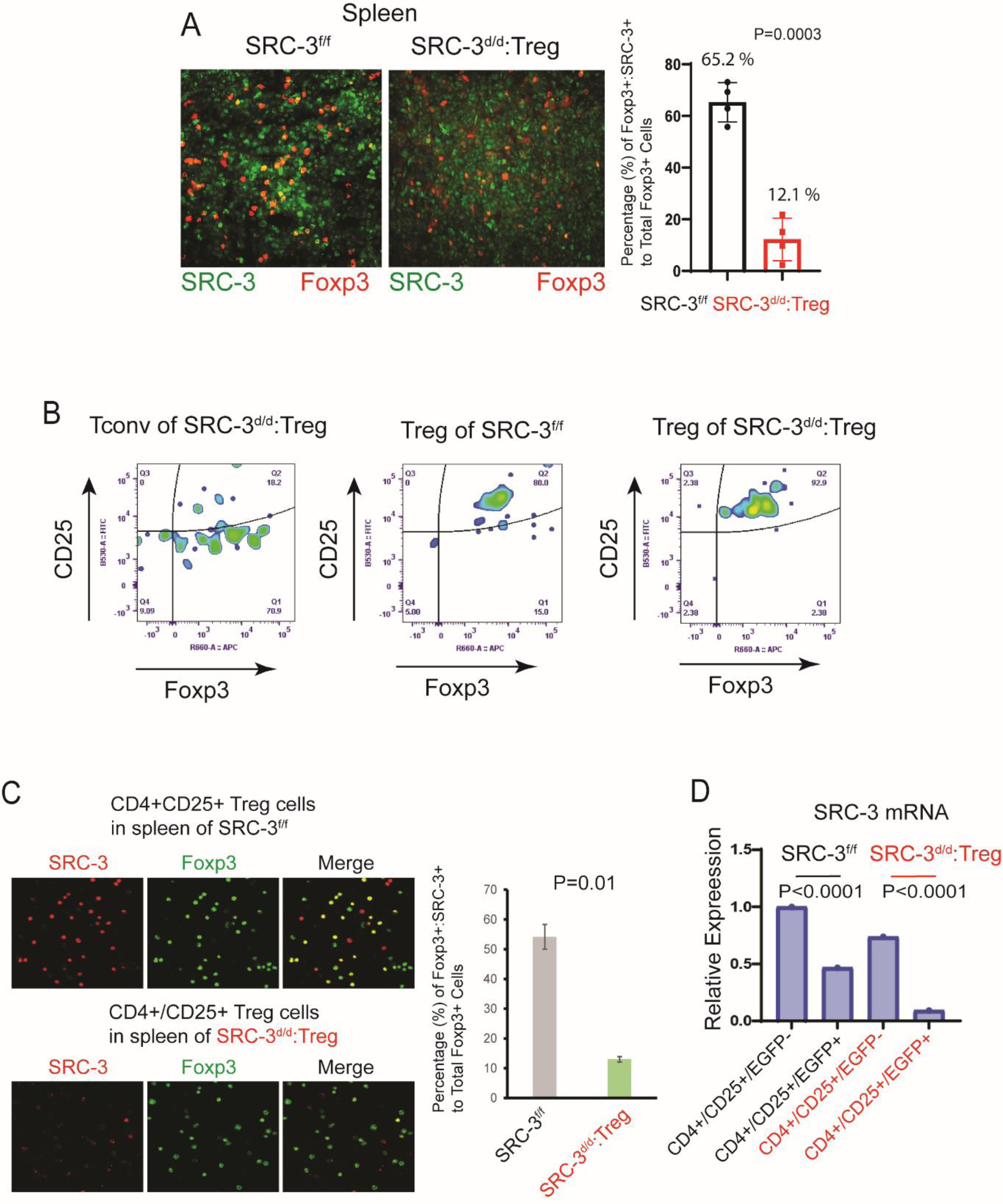
Validation of SRC-3 gene disruption in SRC-3 KO Tregs. (A) Dual immunofluorescence for Foxp3^+^SRC-3^+^ cells in the spleens of SRC-3^f/f^ and SRC-3^d/d^:Treg female mice along with percentage calculations of Foxp3+SRC-3+ cells with respect to the total number of Foxp3+ cells. (B) Validation of isolated Tconv and Tregs from spleens of SRC-3^f/f^ and SRC-3^d/d^:Treg female mice by flow cytometry with antibodies against CD25 and Foxp3. (C) Dual immunofluorescence for Foxp3^+^SRC-3^+^ cells in CD4^+^CD25^+^ Treg cells isolated from spleens of SRC-3^f/f^ and SRC-3^d/d^:Treg female mice along with percentage calculations of Foxp3^+^SRC-3^+^ cells with respect to the total number of Foxp3+ cells. (D) Quantitative PCR (qPCR) for SRC-3 mRNA levels in SRC-3 KO Tregs. CD4^+^CD25^+^EGFP^−^ and CD4^+^CD25^+^EGFP^+^ T cells were isolated from the spleens of SRC-3^f/f^ and SRC-3^d/d^:Treg female mice. RNA was isolated from these T cells, and then SRC-3 mRNA was determined by qPCR with primers corresponding to exon 11 of the SRC-3 mRNA.

**Supplement Fig. S4.**
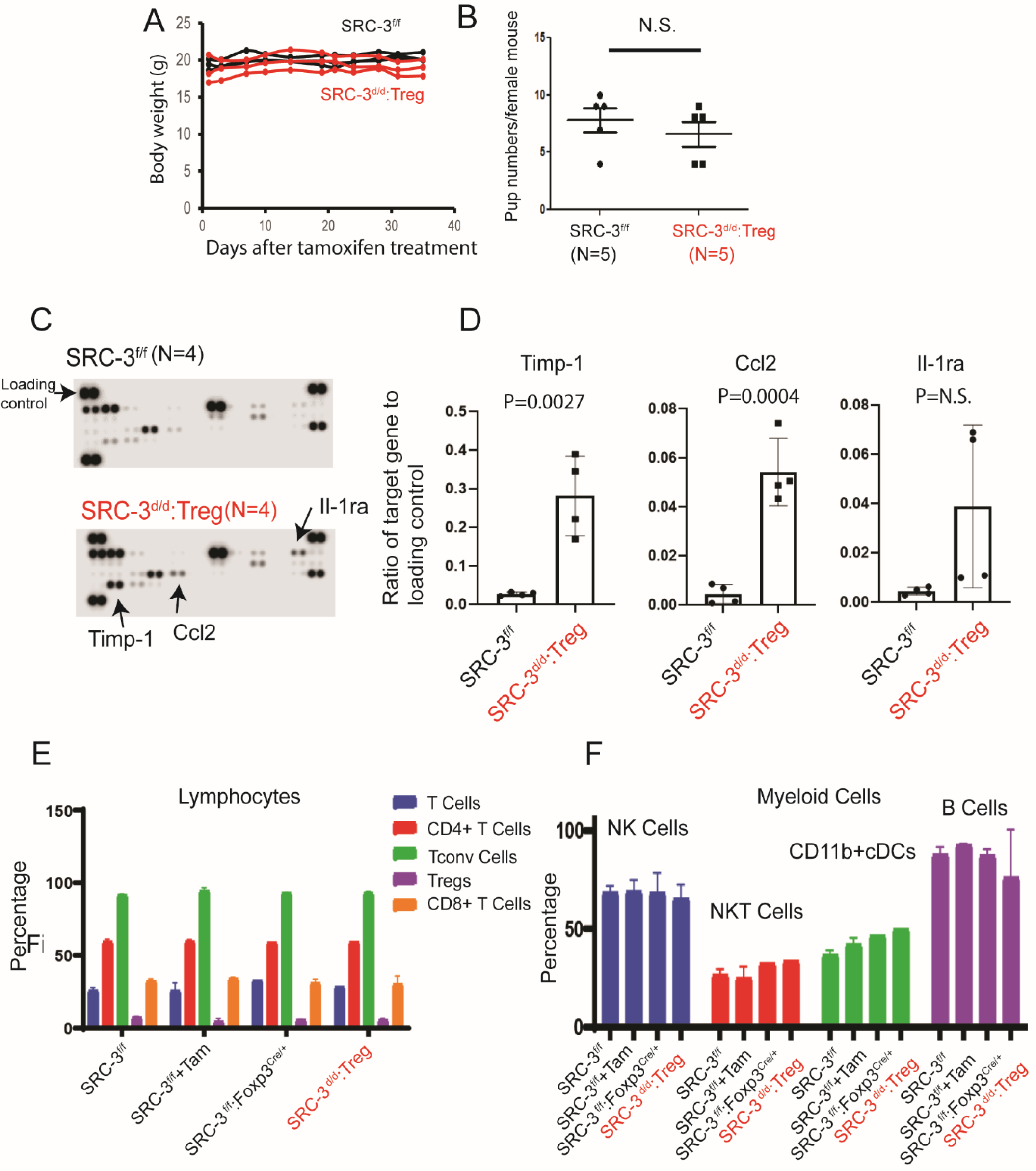
Phenotype of SRC-3d/d:Treg mice. (A) Body weight measurements of SRC-3^f/f^ and SRC-3^d/d^:Treg female mice. These mice were treated with tamoxifen (five days) and then their weight was measured. (B) Fertility of SRC-3^f/f^ versus SRC-3^d/d^:Treg female mice. The number of pups born to SRC-3^f/f^ and SRC-3^d/d^:Treg female mice mated with wild-type male proven-breeders. (C) Cytokine profiles in blood from SRC-3^f/f^ and SRC-3^d/d^:Treg female mice. Blood was collected from these mice on the 30^th^ day after tamoxifen treatment. (D) Quantification of Timp-1, Ccl2, and Il-1ra levels in the blood of SRC-3^f/f^ and SRC-3^d/d^:Treg female mice is shown in panel (C). (E-F) Immunophenotyping of lymphocytes (E) and myeloid cells (F) in spleens from SRC-3^f/f^, tamoxifen-treated SRC-3^f/f^, SRC-3^f/f^:Foxp3^Cre-ERT2/+^, and tamoxifen-treated SRC-3^f/f^:Foxp3^Cre-ERT2/+^ female mice. Spleens were harvested from these mice on the 30^th^ day after either tamoxifen or vehicle treatment. NS, Non Specific.

**Supplement Fig. S5.**
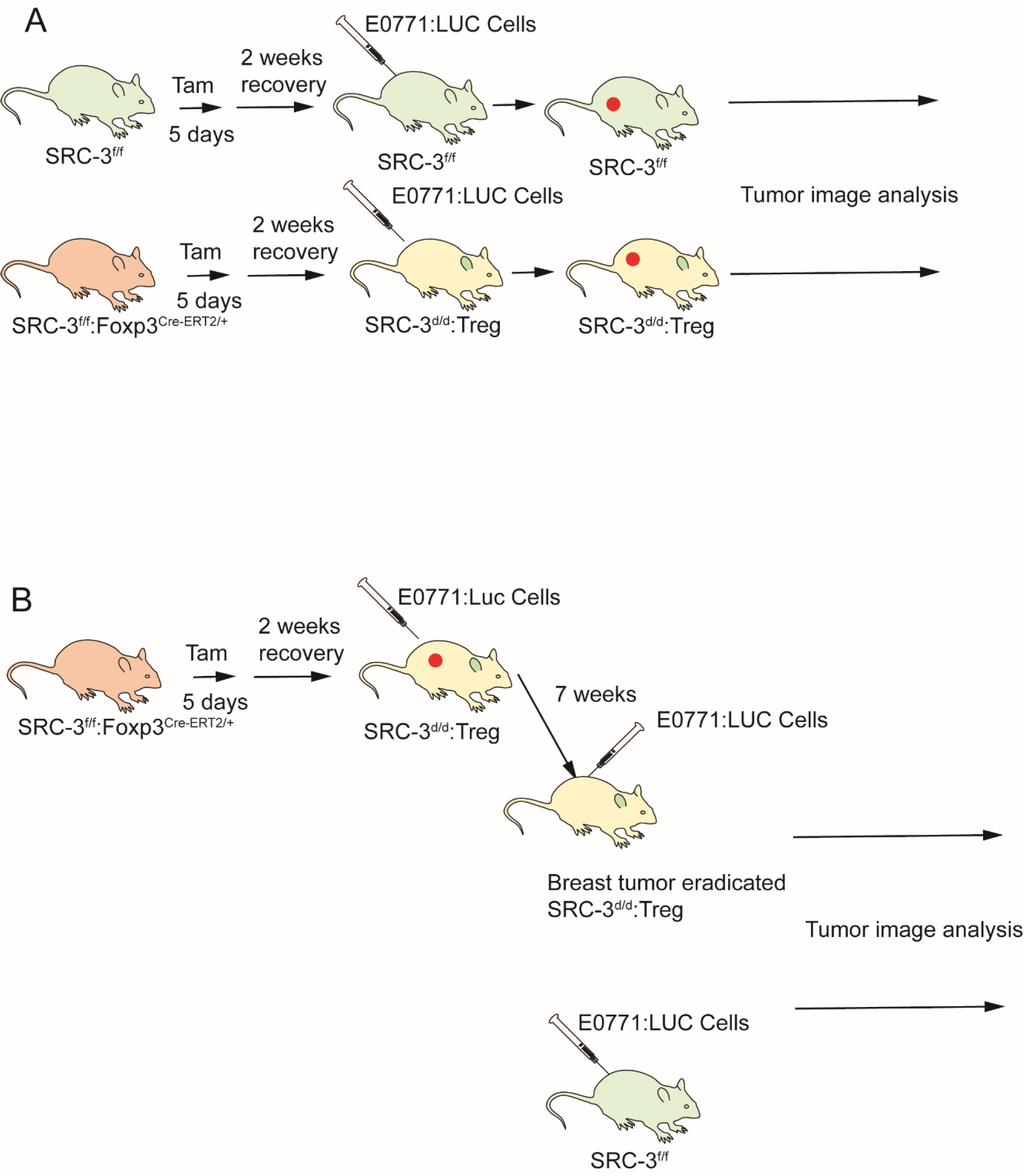
Schematic diagram for SRC-3KO Treg mediated tumor eradication and resistance in SRC-3^d/d^:Treg female mice. (A) Diagram showing tumor eradication in SRC-3^d/d^:Treg female mice. SRC-3^f/f^ and SRC-3^f/f^:Foxp3^Cre-ERT2^ female mice were treated with tamoxifen (75 mg/kg for 5 days). After two weeks, E0771:LUC cells (1×10^5^ cells) were orthotopically injected into the mammary fat pad of SRC-3^f/f^ and SRC-3^d/d^:Treg female mice. Afterward, tumor luciferase activity images were collected. (B) Diagram defining tumor resistance in SRC-3^d/d^:Treg female mice. E0771:LUC cells were orthotopically injected into SRC-3^d/d^:Treg mice. After seven weeks, tumors were eradicated in SRC-3^d/d^:Treg female mice, and then mice were orthotopically reinjected with E0771:LUC cells. As controls, E0771 cells were orthotopically injected into SRC-3^f/f^ female mice.

**Supplement Fig. S6.**
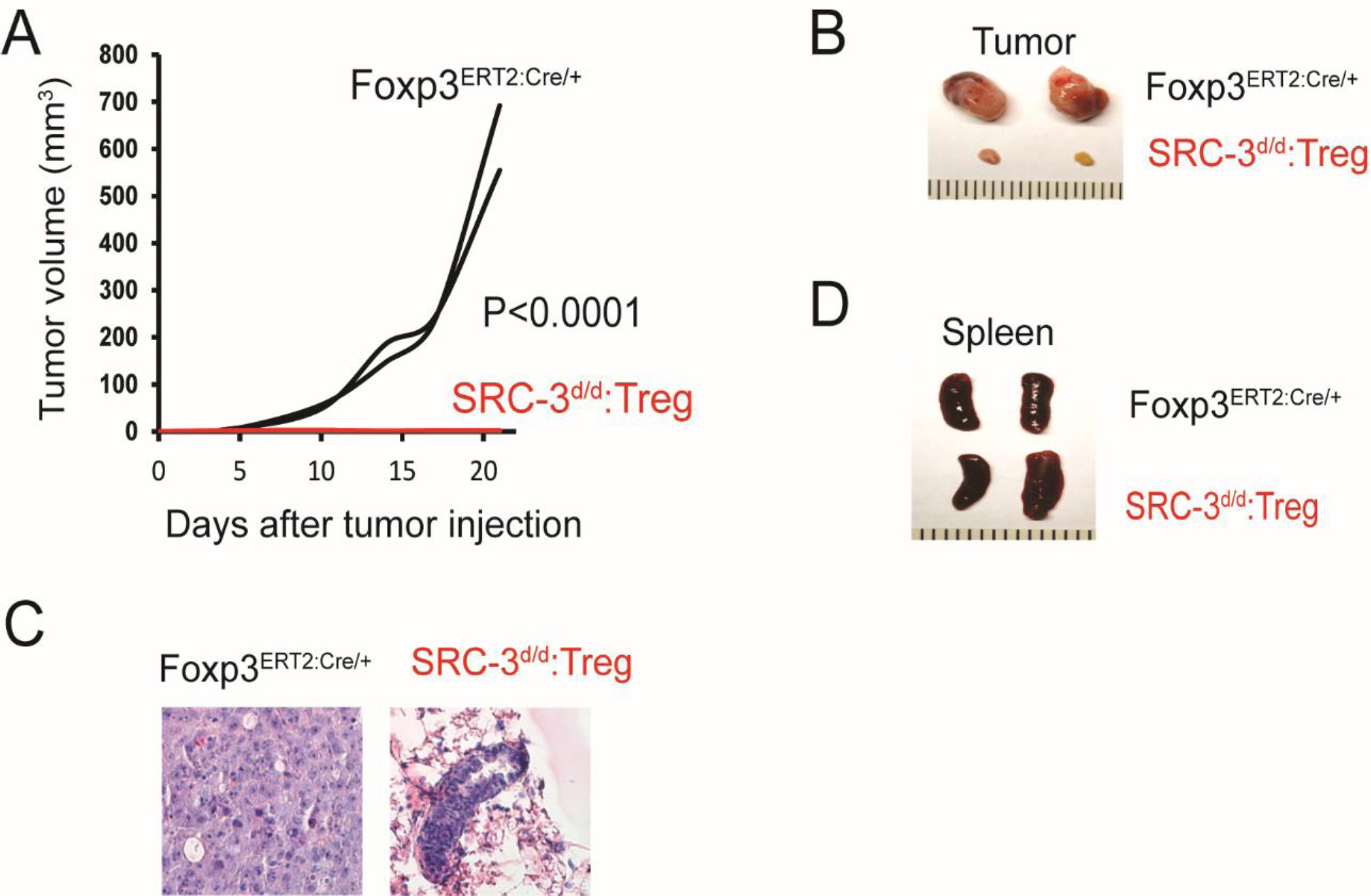
Growth of E0771 breast tumors in Foxp3Cre-ERT2 female mice. (A) E0771 breast tumor growth in Foxp3^Cre-ERT2^ and SRC-3^d/d^:Treg female mice. (B) Breast tumors isolated from Foxp3^Cre-ERT2^ and SRC-3^d/d^:Treg female mice on the 23^rd^ day after injection of E0771 cells. (C) H&E staining of tumors is presented in panel (B). (D) Spleens isolated from Foxp3^Cre-ER^ and SRC-3^d/d^:Teg female mice on the 40^th^ day after tamoxifen treatment.

**Supplement Fig. S7.**
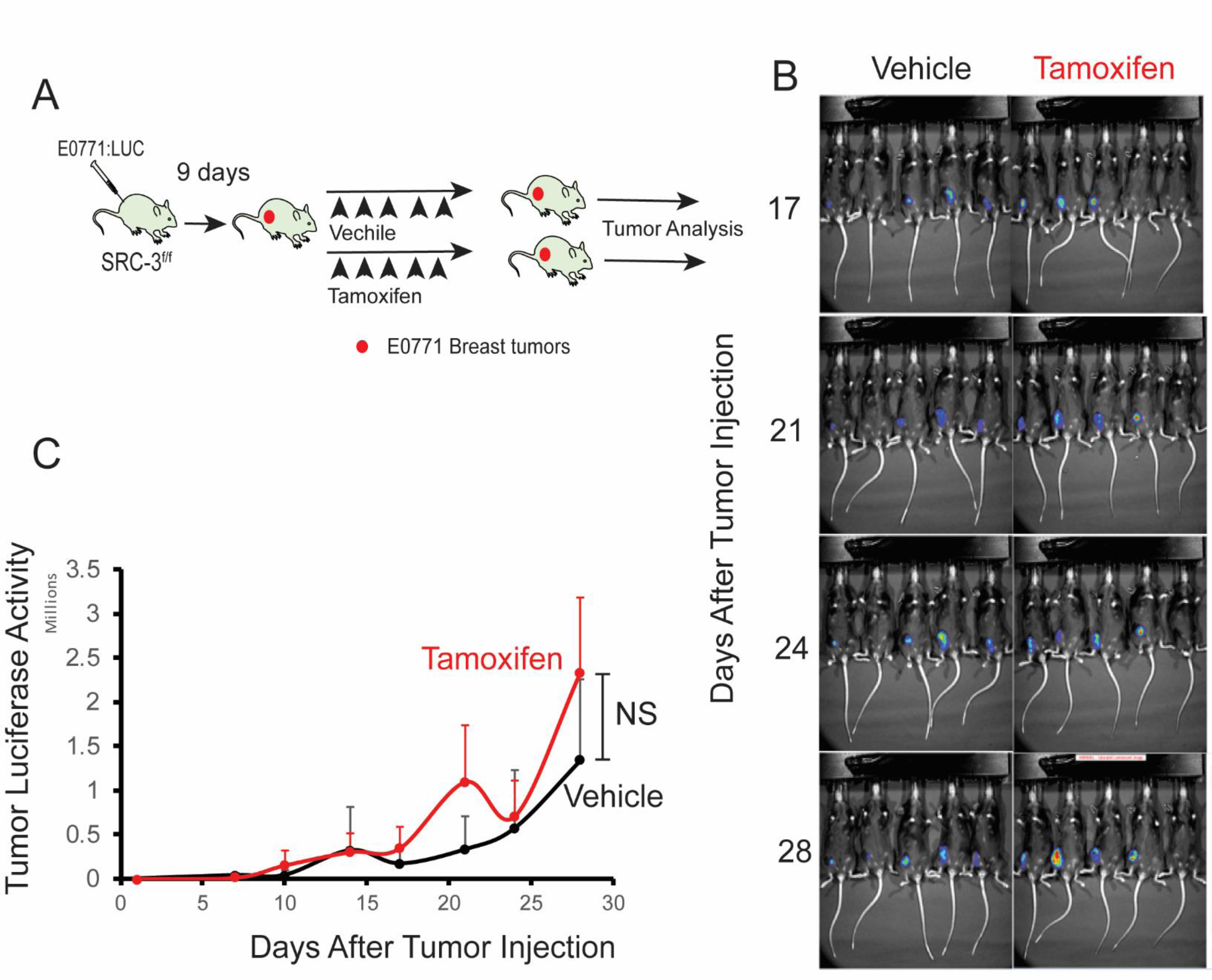
Tamoxifen treatment did not impact the growth of E0771 breast tumors in SRC-3^f/f^ female mice. (A) Schematic describing tamoxifen treatment in tumor-bearing SRC-3^f/f^ female mice. (B) Tumor luciferase activity of tumor-bearing SRC-3^f/f^ female mice treated with tamoxifen versus vehicle. (C) The quantification of tumor luciferase activity in Panel B.

**Supplement Fig. S8.**
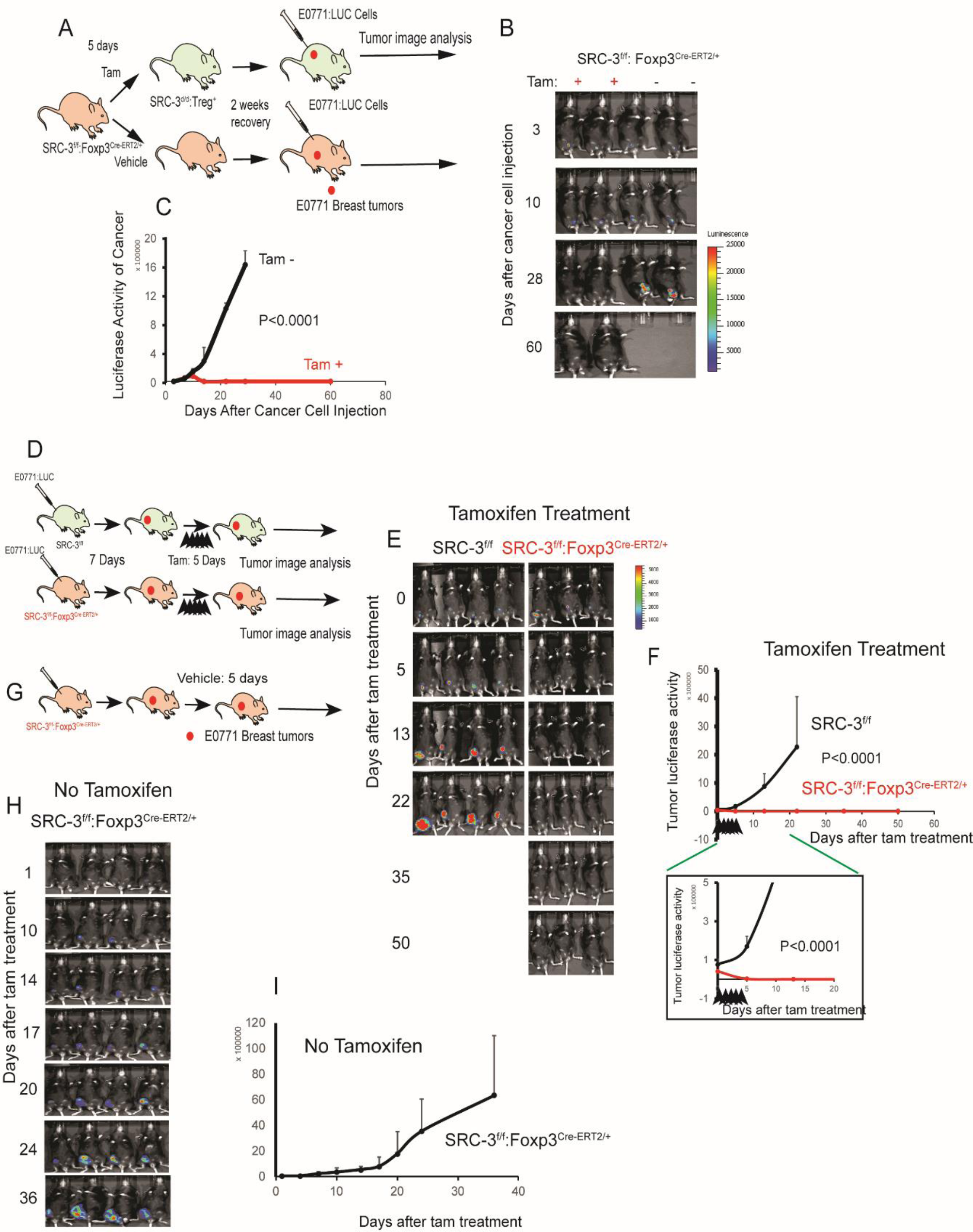
Breast tumor regression and prevention in SRC-3d/d:Treg female mice. (A) Diagram to illustrate the suppression of tumor initiation by SRC-3 KO Treg cell induction. SRC-3^f/f^:Foxp3^Cre-ERT2/+^ female mice were treated with tamoxifen to generate SRC-3^d/d^:Treg mice. As controls, SRC-3^f/f^:Foxp3^Cre-ERT2/+^ female mice were treated with vehicle instead. E0771 cells (1×10^5^ cells) were orthotopically injected into SRC-3^f/f^:Foxp3^Cre-ERT2/+^ and SRC-3^d/d^:Treg mice two weeks later after tamoxifen treatment. (B) Tumor luciferase activity images are presented in panel (A). (C) Quantification of the tumor luciferase images are presented in panel (B). (D) The diagram illustrates the tumor regression activity of SRC-3 KO Tregs. SRC-3^f/f^ and SRC-3^f/f^:Foxp3^Cre-ERT2/+^ female mice were orthotopically injected with E0771 cells. After seven days, tumor-bearing SRC-3^f/f^ and SRC-3^f/f^:Foxp3^Cre-ERT2/+^ mice were treated with tamoxifen for five days. Afterward, the luciferase activity emanating from tumors was determined. (E) Determination of luciferase activity from tumors is presented in panel (D). (F) Quantifying luciferase activity is presented in panel **e** until the ^2^0th day, which is in the box, and the 50^th^ day after tamoxifen treatment. (G) This diagram illustrates the tumor progression in SRC-3^f/f^:Foxp3^Cre-ERT2/+^ female mice upon vehicle treatment. (H) The tumor luciferase activity in panel G. (I) Quantitation of luciferase activity is presented in panel H. Tam, Tamoxifen.

**Supplement Fig. S9.**
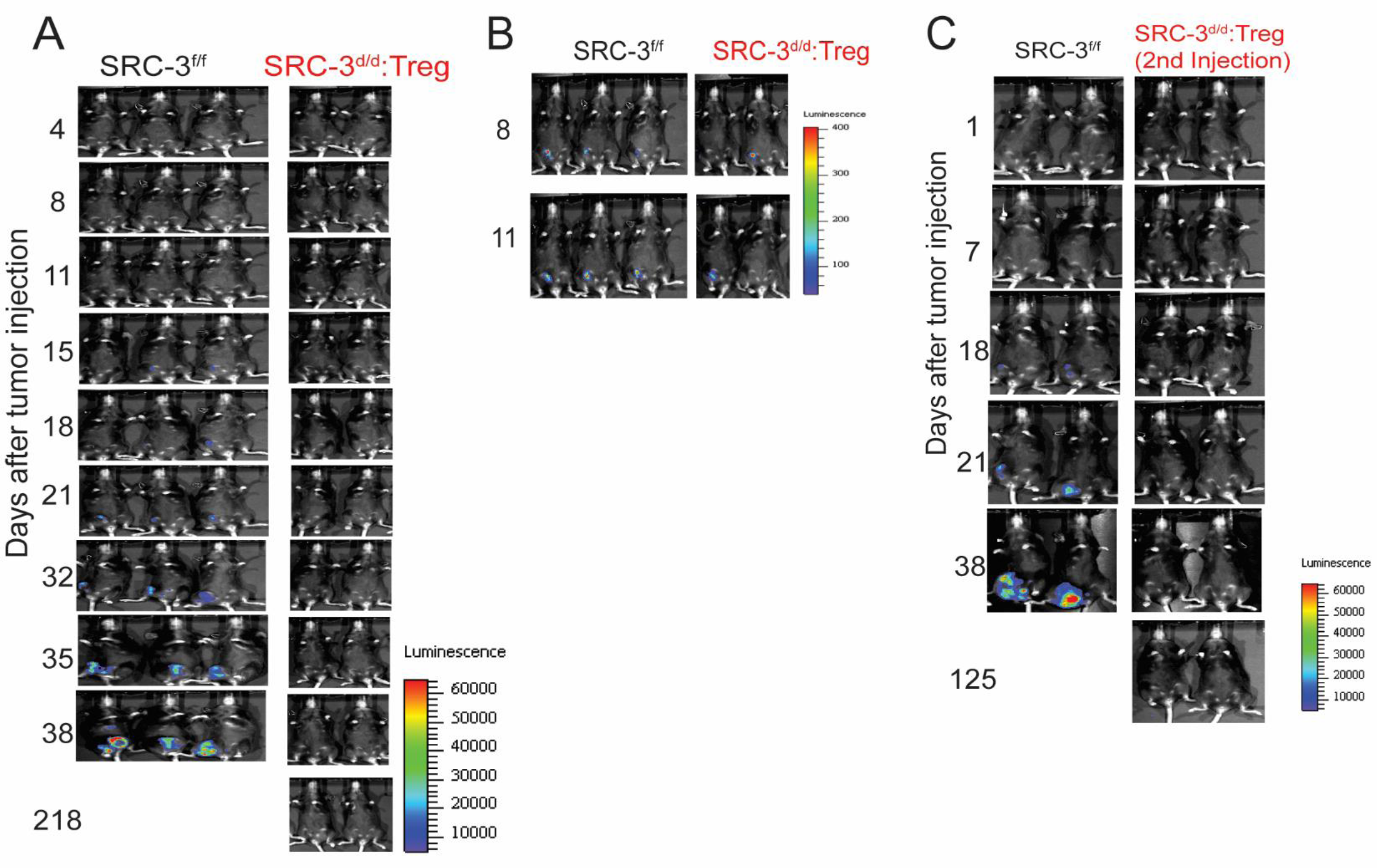
E0771 breast tumor eradication in SRC-3^d/d^:Treg mice. (A) Breast tumor luciferase activity in SRC-3^f/f^ and SRC-3^d/d^:Treg female mice. (B) Tumor luciferase activity in tamoxifen-treated SRC-3^f/f^ and SRC-3^d/d^:Treg mice on the 8^th^ and 11^th^ days after E0771 cell injection. (C) Tumor luciferase activity in SRC-3^f/f^ and breast cancer-eradicated SRC-3^d/d^:Treg mice after a 2^nd^ injection of E0771:LUC cells.

**Supplement Fig. S10.**
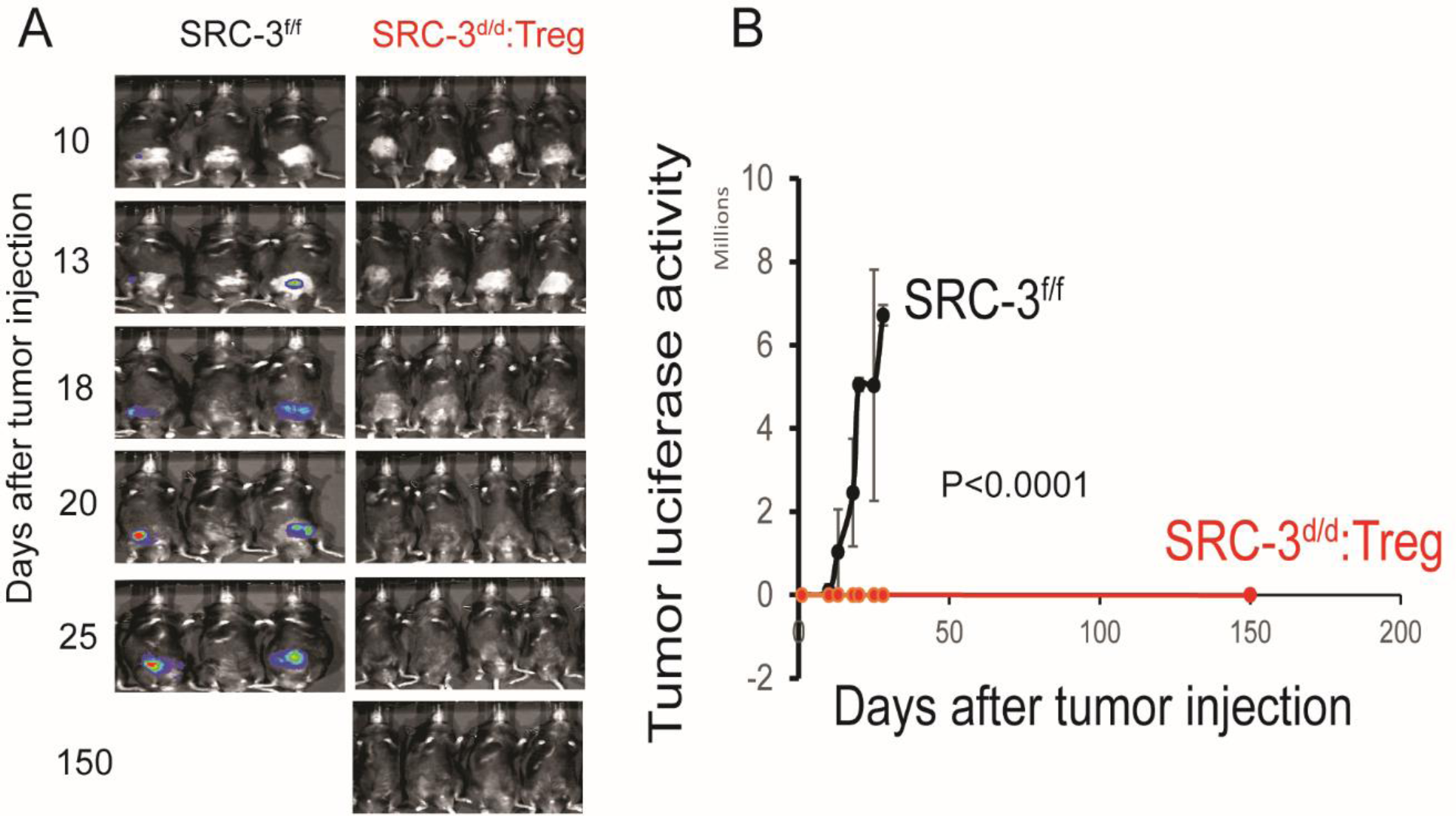
Eradication of prostate cancer in SRC-3^d/d^:Treg male mice. A) Determination of the luciferase activity emanating from prostate tumors in SRC-3^f/f^ and SRC-3^d/d^:Treg male mice. Luciferase-labeled RM-1 mouse prostate cancer cells (1×10^3^ cells) were orthotopically injected into the prostate of SRC-3^f/f^ and SRC-3^d/d^:Treg male mice. Afterward, tumor luciferase activity arising from the prostate was determined. (B) Quantification of prostate tumor luciferase activity is presented in panel (A).

**Supplement Fig. S11.**
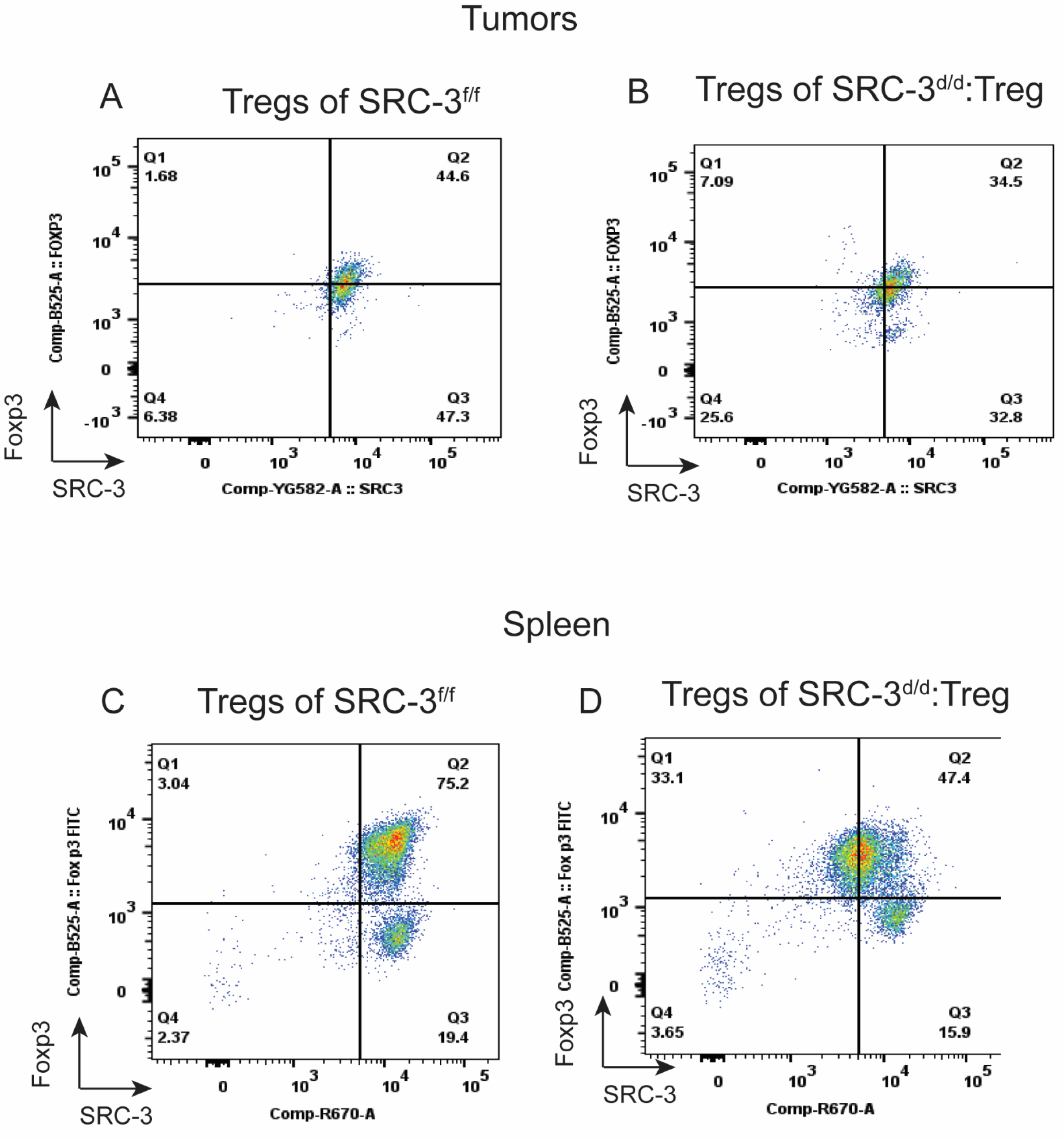
Relative number of SRC-3 KO Tregs in tumors and spleens. (A and B) The pattern of SRC-3 and Foxp3 expression in tumor Tregs from SRC-3^f/f^ (A) and SRC-3^d/d^:Treg male mice (B) determined by flow cytometry with antibodies against SRC-3 and Foxp3. (C and D) The pattern of SRC-3 and Foxp3 levels in spleen Tregs from SRC-3^f/f^ (C) and SRC-3^d/d^:Treg male mice (D) determined by flow cytometry with antibodies against SRC-3 and Foxp3.

**Supplement Fig. S12.**
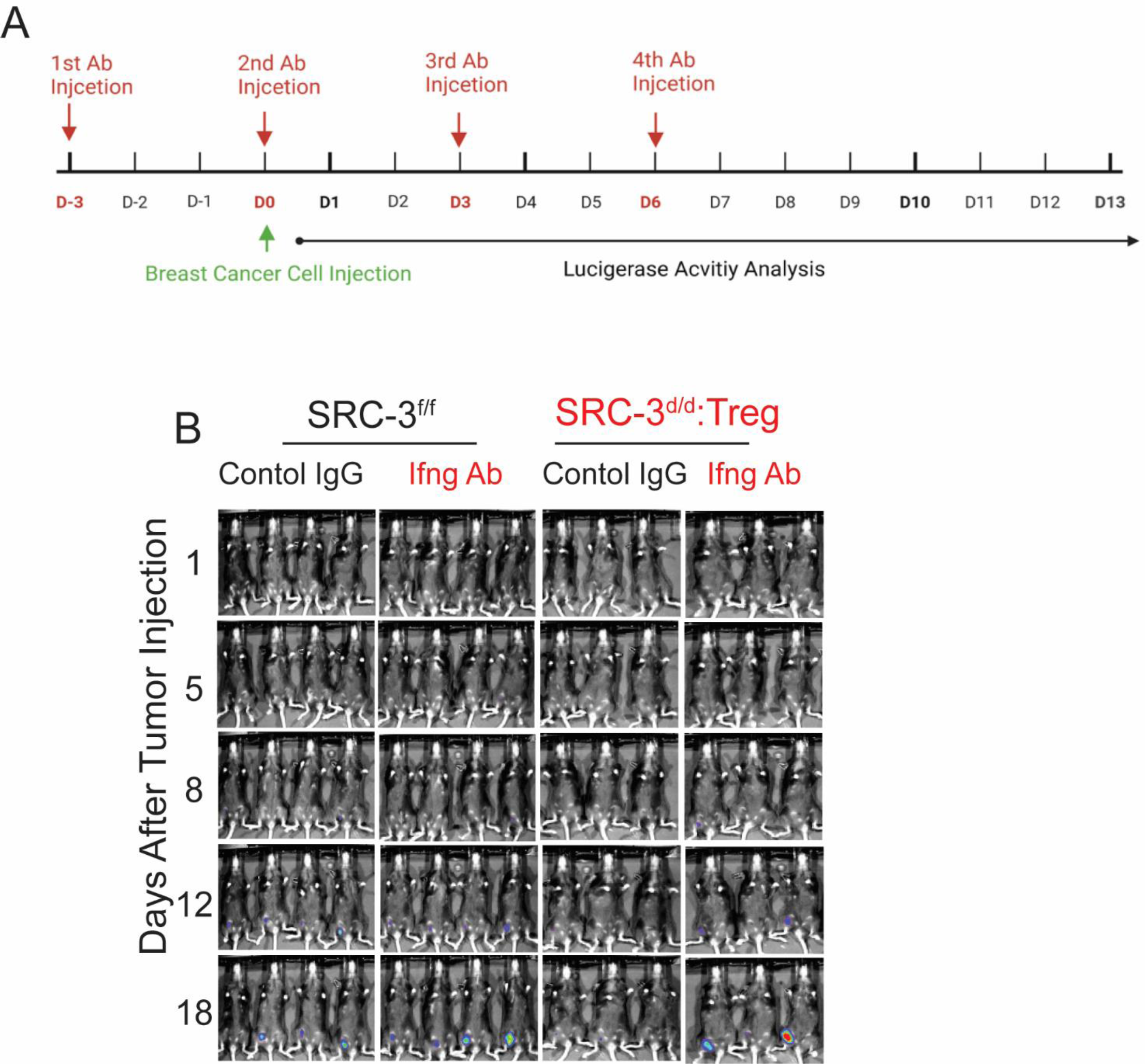
Anti-Ifng antibody treatment prevents tumor eradication activity of SRC-3^d/d^ :Foxp3^Cre/+^ Tregs in female mice. (A) Schematic for anti-Ifng antibody and control rat IgG treatment of breast tumor-bearing SRC-3^f/f^ and SRC-3^d/d^:Treg mice. (B) Tumor luciferase activity of tumor-bearing SRC-3^f/f^ and SRC-3^d/d^:Treg female mice treated with anti-Ifng antibody (Ifng Ab) and control rat IgG (Control IgG).

**Supplement Fig. S13.**
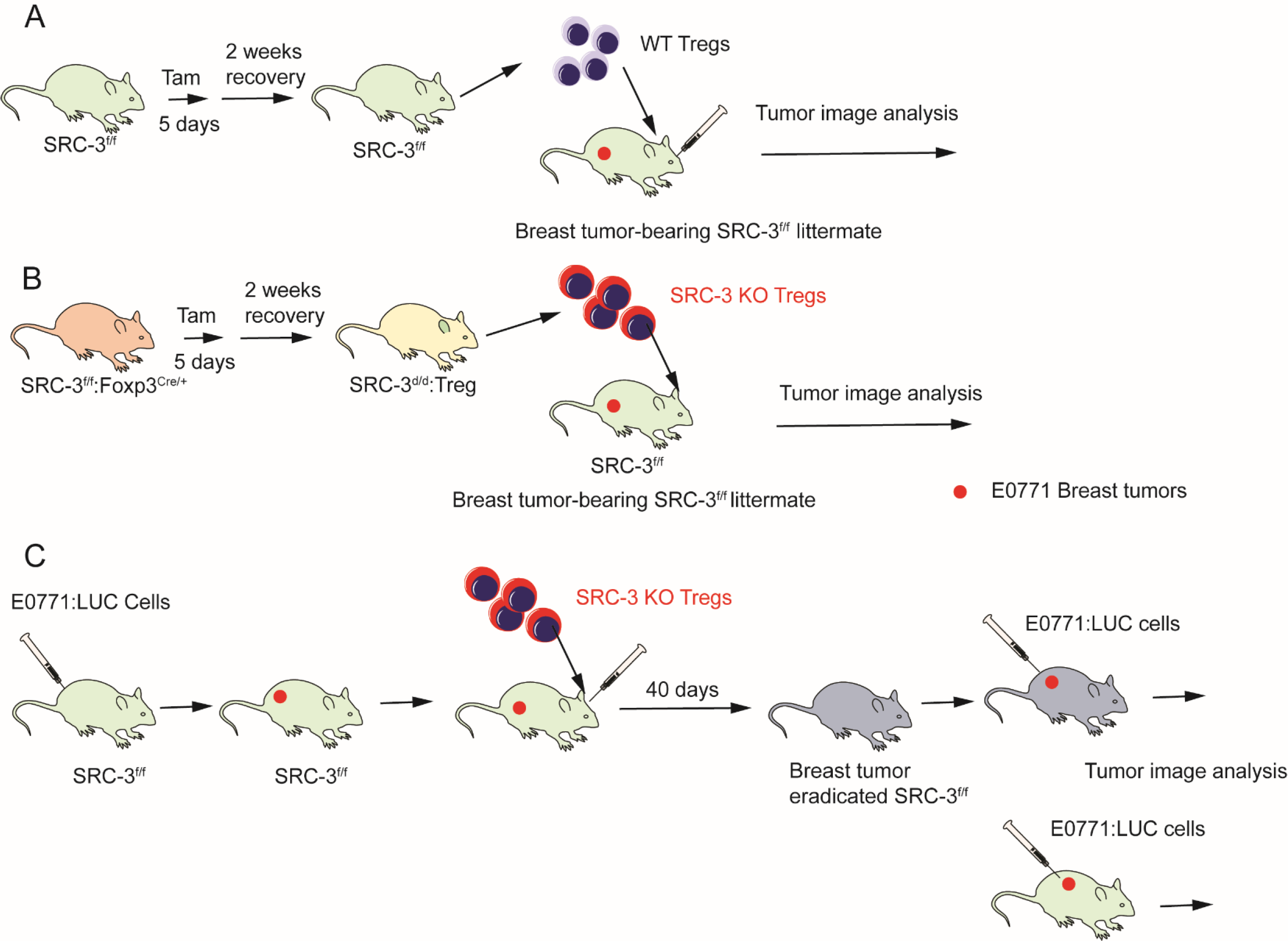
Schematic diagram of the SRC-3KO Treg ACT process. (A) ACT with wild-type Tregs was performed by isolating Tregs from the spleen of SRC-3^f/f^ female mice treated with tamoxifen. These Tregs were then administrated into tumor-bearing SRC-3^f/f^ female littermates by retro-orbital injection. (B) ACT with SRC-3 KO Tregs was performed similarly, except that SRC-3^d/d^:Treg female mice were used as Treg donors. (C) Tumor-resistant mice generated after ACT with SRC-3 KO Tregs. SRC-3 KO Tregs were isolated from the spleens of SRC-3^d/d^:Treg female mice. Tumor-bearing SRC-3^f/f^ female mice were adoptively transferred with SRC-3 KO Tregs. Tumors were eradicated in these SRC-3^f/f^ female mice by the 40^th^ day after ACT. Afterward, E0771:LUC cells (1×10^5^ cells) were orthotopically injected into tumor-eradicated SRC-3^f/f^ female mice adoptively transferred with SRC-3 KO Tregs. As controls, E0771:LUC cells (1×10^5^ cells) were orthotopically injected into SRC-3^f/f^ female mice. Tam, Tamoxifen.

**Supplement Fig S14.**
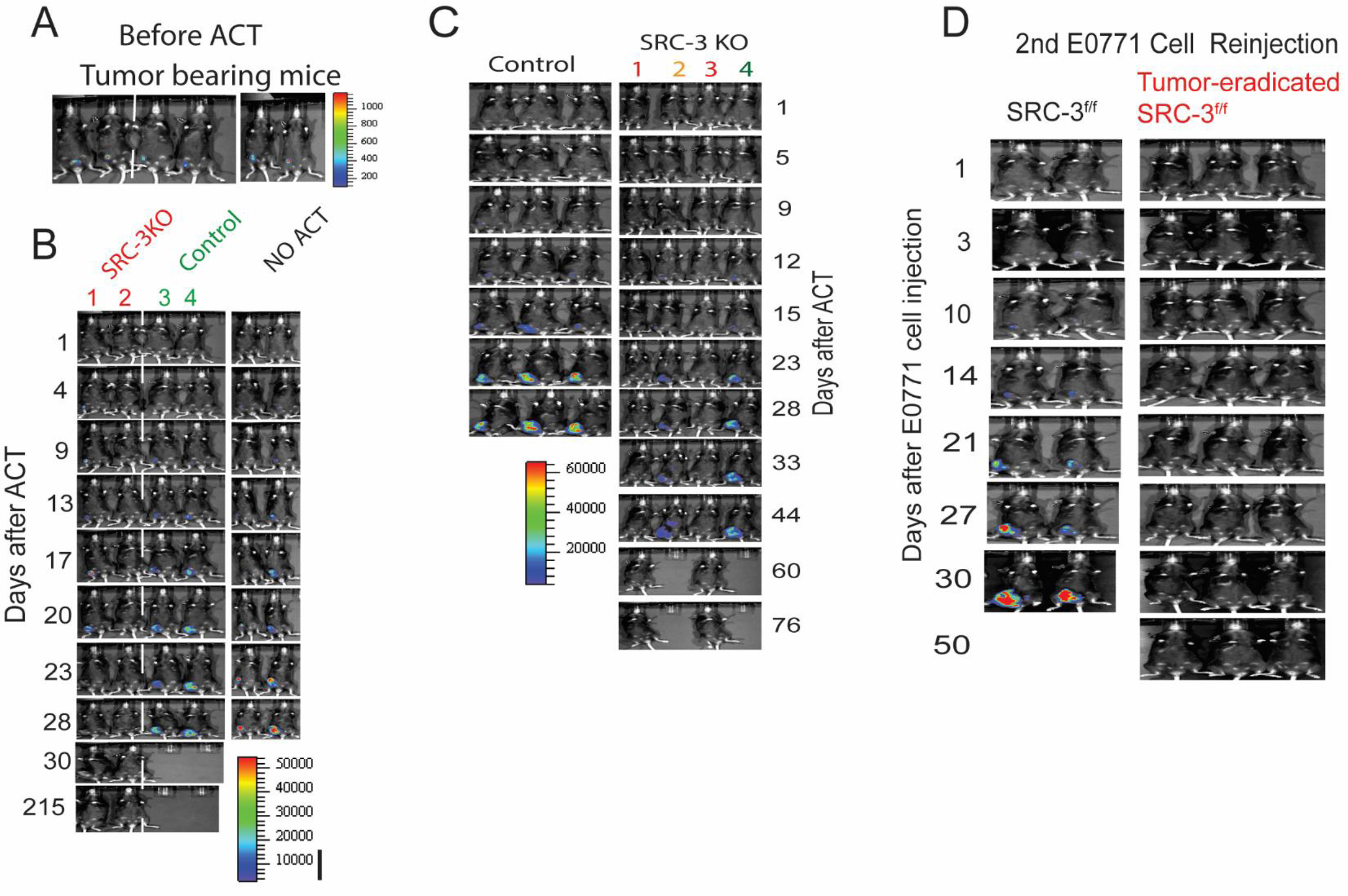
Tumor eradication through adoptive SRC-3 KO Treg transfer. (A) Tumor luciferase activity in tumor-bearing SRC-3^f/f^ female mice before ACT. (B) Tumor luciferase image analysis in tumor-bearing SRC-3^f/f^ female mice after ACT with SRC-3 KO Tregs (800K cells) and wild-type Tregs (800K cells). Control tumor-bearing SRC-3^f/f^ mice that did not undergo ACT (NO ACT). (C) Tumor luciferase activity in SRC-3^f/f^ female mice after ACT with different doses of SRC-3 KO Tregs. Tumor-bearing SRC-3^f/f^ female mice were injected with adoptively transferred wild-type (800K cells) or SRC-3 KO Treg cells (1:610 K cells, 2:326 K cells, 3:643 K cells, 4:320 K cells). Afterward, the luciferase activity emanating from tumors was determined. (D) Tumor luciferase activity in tumor-eradicated SRC-3^f/f^ female mice after ACT with SRC-3 KO Tregs and SRC-3^f/f^ female control mice after a 2^nd^ injection of E0771:LUC cells.

**Supplement Fig. S15.**
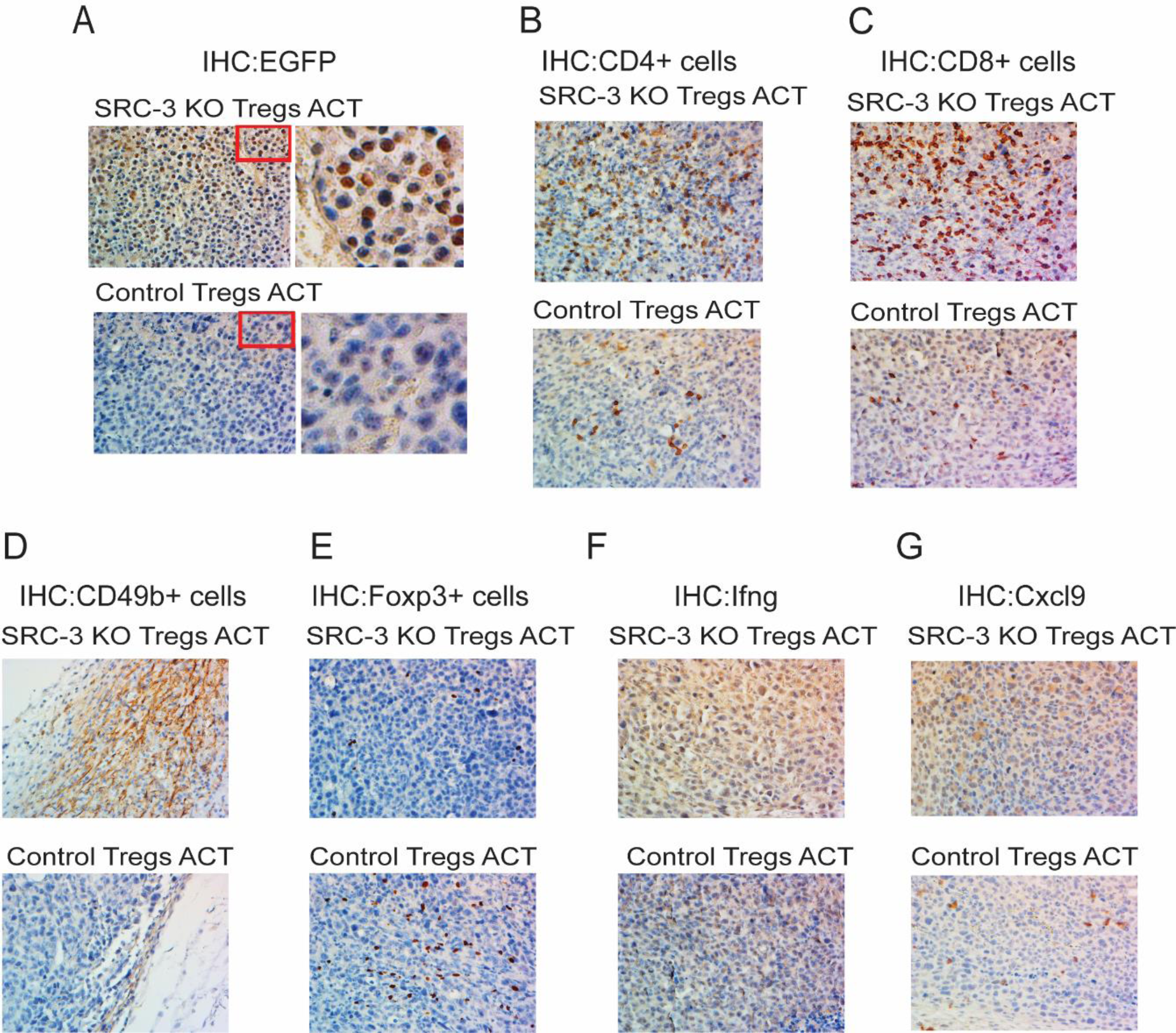
Breast tumors acquire an antitumor immune environment after ACT with SRC-3 KO Tregs. (A-G) IHC analysis of expression of EGFP (A), CD4+ (B), CD8+(C), CD56+ (D), Foxp3 (E), Ifng (F) and Cxcl9 (G) in breast tumors in SRC-3^f/f^ female mice receiving ACT with SRC-3 KO or WT control Tregs.

**Supplement Fig. S16.**
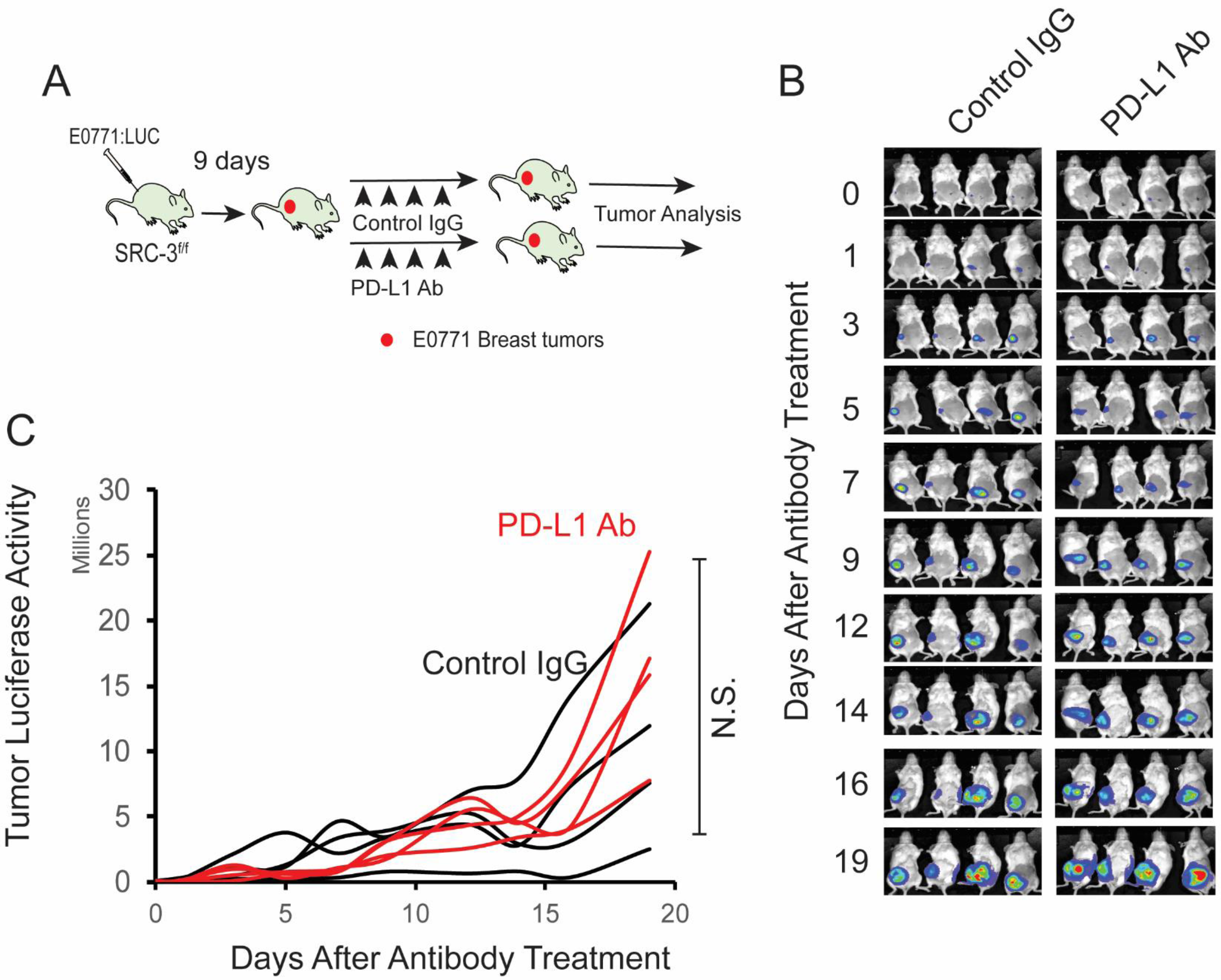
Anti PD-L1 antibody did not suppress E0771 breast tumor progression in SRC-3^f/f^ female mice. (A) Schematic for anti-PD-L1 antibody treatment of breast tumor-bearing SRC-3^f/f^ female mice. (B) Tumor luciferase activity of tumor-bearing SRC-3 ^f/f^ female mice treated with anti-PD-L1 antibody (PD-L1 Ab, 100 μg/20g mouse) and control rat IgG (Control IgG, 100 μg/20g mouse). (C) The quantification of tumor luciferase activity in Panel B.

---

**Supplement Table 1.**
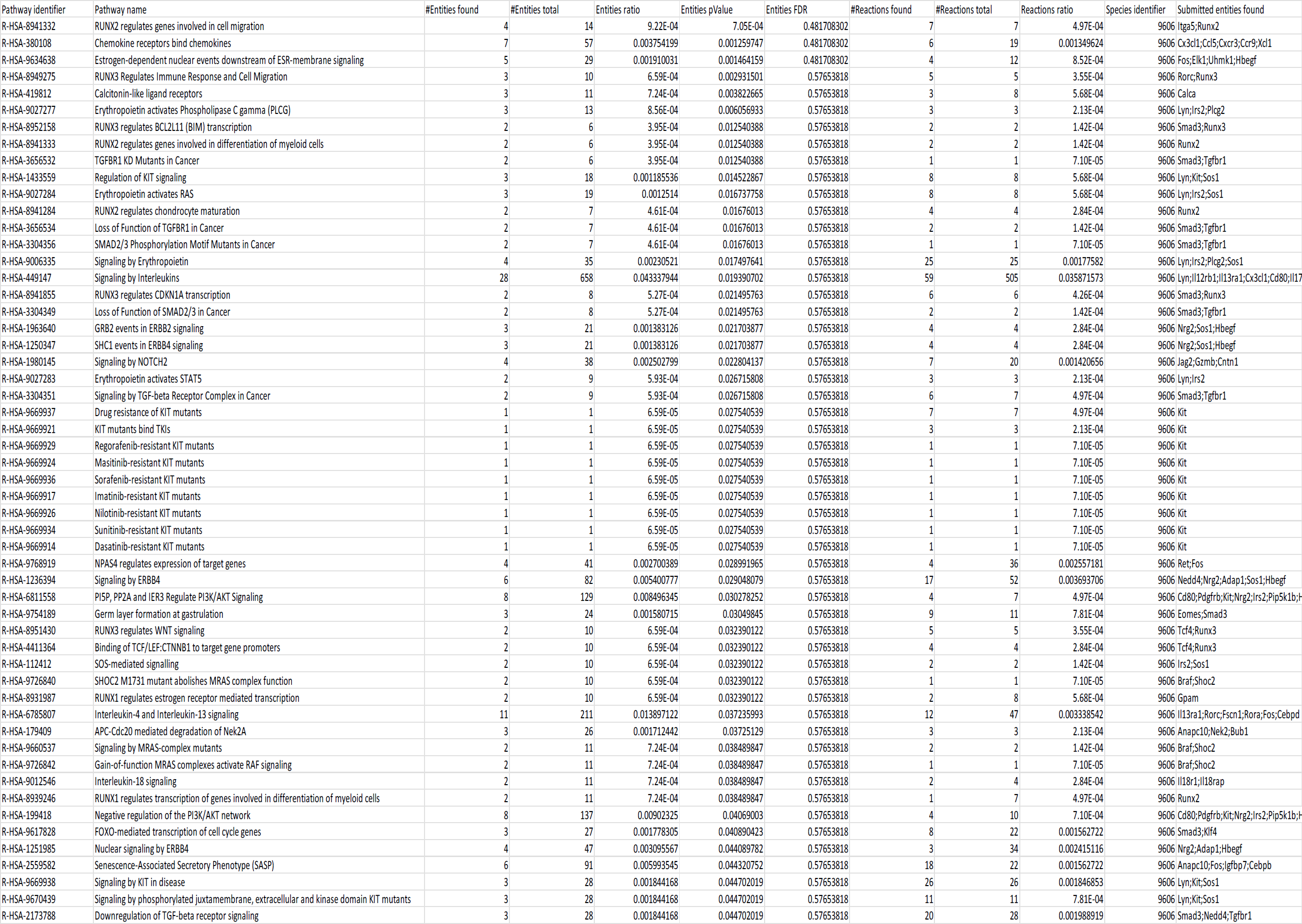
Downregulated pathways in SRC-3 KO Treg cells.

**Supplement Table 2.**
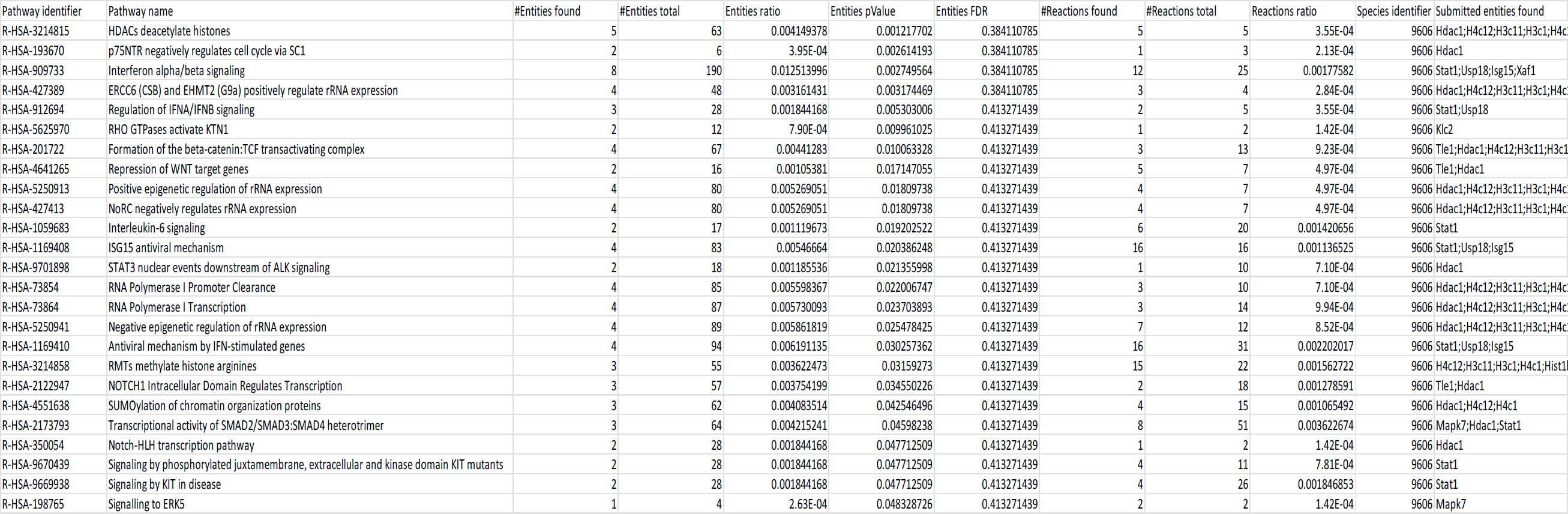
Upregulated pathways in SRC-3 KO Treg cells.

